# Genetically engineered ESC-derived embryos reveal Vinculin-dependent force responses required for mammalian neural tube closure

**DOI:** 10.64898/2025.12.22.696028

**Authors:** Ian S. Prudhomme, Eric R. Brooks, Nilay Taneja, Bhaswati Bhattacharya, Brian J. LaFleche, Yasuhide Furuta, Jennifer A. Zallen

## Abstract

Epithelial sheets build complex structures by converting mechanical forces into changes in cell and tissue organization. During neural tube closure, the neural plate dynamically remodels to produce a closed tube that provides the structural foundation for the developing brain and spinal cord. How cells maintain epithelial integrity despite the forces required for tissue morphogenesis during neural tube closure is not understood. We show that mechanical forces are upregulated during cranial neural tube closure in the mouse embryo and recruit the force-sensitive protein Vinculin to adherens junctions. Leveraging a genetically engineered embryonic stem cell-based pipeline to efficiently generate mutant embryos, we show that *Vinculin* mutants produce mechanical forces correctly but fail to maintain cell adhesion under tension, resulting in a failure of cranial neural fold elevation. Live imaging of cell behavior in the developing midbrain reveals that apical constriction, cell rearrangement, and cell division initiate correctly in *Vinculin* mutants, but their progression is impeded by disruption of adherens junctions at sites of increased tension. These results demonstrate that Vinculin is required to reinforce cell adhesion in response to increasing physiological forces during cranial neural tube closure, and that this activity is necessary to translate these forces into changes in tissue structure.

## Introduction

The development of multicellular organisms requires the generation of complex three-dimensional structures from simple sheets of cells. Formation of the mammalian brain involves dynamic changes in cell shape and organization that transform an epithelial sheet into the closed neural tube that is essential for the development of the brain and spinal cord (Wilde et al., 2014; Juriloff and Harris, 2018). Disruptions in neural tube closure are among the most common causes of birth defects, affecting around 1 in 2,000 births, representing a significant challenge in human health (Wallingford et al., 2013; Zaganjor et al., 2016). Neural tube closure requires the integration of multiple cell behaviors, including apical constriction, cell division, and cell rearrangements, which promote epithelial reorganization, bending, and fusion (Nikolopoulou et al., 2017; Vijayraghavan and Davidson, 2017; Matsuda and Sokol, 2021). These processes are driven by spatiotemporally regulated actomyosin networks that are predicted to alter the physical environment experienced by individual cells and create mechanical stresses that could disrupt cell adhesion and epithelial integrity. However, the mechanical forces that drive mammalian neural tube closure, and how cells maintain tissue integrity in the presence of these forces, are not understood.

Biophysical studies have identified a number of force-induced changes in protein conformation and activity that could allow cells to maintain cell adhesion and tissue structure under tension. Mechanical forces promote structural changes in E-cadherin, α-catenin, and filamentous actin (F-actin) that are predicted to stabilize protein interactions within junctional complexes and strengthen connections between adherens junctions and the actin cytoskeleton (Yonemura et al., 2010; Rakshit et al., 2012; Buckley et al., 2014; Huang et al., 2017; Sun et al., 2020). In addition, mechanical forces modulate interactions between subunits of cell-matrix adhesions (Burridge and Guilluy, 2016) and epithelial tight junctions (Spadaro et al., 2017; Citi 2019; Citi et al., 2024) that could provide additional strategies to reinforce cell-matrix anchoring and epithelial barrier function. Studies of epithelial remodeling *in vivo* have identified tricellular junctions—where three cells meet at a single point or vertex—as sites of increased tension that engage a variety of force responses in epithelial cells (Higashi and Miller, 2017; Bosveld et al., 2018). Tricellular adherens junctions recruit several proteins that are required to stabilize cell adhesion under tension, both in culture and in epithelial tissues *in vivo* (le Duc et al., 2010; Sawyer et al., 2011; Thomas et al., 2013; Choi et al., 2016; Razzell et al., 2018; Rauskolb et al., 2019; Yu and Zallen, 2020; Perez-Vale et al., 2021; Cavanaugh et al., 2022; Sheppard et al., 2023). In addition, components of tricellular tight junctions are required to maintain epithelial barrier function under tension in cultured cells and in the *Xenopus* embryo (Oda et al., 2014; Stephenson et al., 2019; Cho et al., 2022; van den Goor et al., 2024). However, how epithelial cells maintain cell adhesion at tricellular junctions in the presence of the physiological forces required to drive epithelial remodeling during mammalian embryogenesis is poorly understood, in part due to the challenges of directly visualizing dynamic processes in mutant embryos.

An excellent candidate for mediating force responses in mammalian tissues is the junction-actin linker protein Vinculin. Vinculin has been shown to be recruited by mechanical forces to cell-matrix adhesions, adherens junctions, and tight junctions across a broad array of cell types (Goldmann, 2016; Bays and DeMali, 2017; Citi, 2019), and is required to reinforce connections between cell adhesion complexes and the actin cytoskeleton in response to mechanical forces *in vitro* (Maddugoda et al., 2007; Peng et al., 2010; le Duc et al., 2010; Yonemura et al., 2010; Taguchi et al., 2011; Huveneers et al., 2012; Peng et al., 2012; Huang et al., 2017). In addition to linking junctions to actin, Vinculin also regulates myosin II activity and actin organization at adherens junctions through interactions with actin regulatory proteins (Wen et al., 2009; Le Clainche et al., 2010; le Duc et al., 2010; Leerberg et al., 2014; Ito et al., 2017). Vinculin has essential *in vivo* roles in regulating heart development and function (Xu et al., 1998; Zemljic-Harpf et al., 2007; Cheng et al., 2016; Fukuda et al., 2019), vascular remodeling (Carvalho et al., 2019; Kotini et al., 2022), stem cell proliferation (Biswas et al., 2021; Bohere et al, 2022), and epithelial and endothelial barrier function (Higashi et al., 2016; van der Stoel et al., 2023; van den Goor et al., 2024; Landino et al., 2025). Furthermore, mouse embryos lacking *Vinculin* display a fully penetrant failure of cranial neural tube closure (Xu et al., 1998; Marg et al., 2010) and mutations in human Vinculin are associated with increased risk of neural tube defects (Wang et al., 2021). However, the mechanisms by which Vinculin regulates mammalian neural tube closure, and how Vinculin influences junctional organization and cell behavior during this process, are unknown.

Here we used a high-throughput embryonic stem cell-based method to efficiently generate large numbers of mutant embryos in order to analyze the role of Vinculin during mouse cranial neural tube closure. Using biophysical and live imaging approaches, we show that mechanical forces in the developing midbrain are anisotropic and increase as neural tube closure proceeds. Mechanical forces are generated correctly in *Vinculin* mutants, but neuroepithelial cells fail to maintain cell adhesion at high-tension junctions as forces increase during neural fold elevation, resulting in a failure of cranial neural tube closure. The defects in *Vinculin* mutants were most pronounced at adherens junctions, whereas tight junctions remained largely intact, and resulted in a progressive disruption of cell adhesion, cell division, and apical constriction behaviors during cranial neural tube closure. These results demonstrate that Vinculin is necessary to stabilize cell adhesion in the presence of endogenous forces in the mouse cranial neural plate and suggest that this function is necessary to convert mechanical forces into changes in cranial neural tube structure.

## Results

### Mechanical forces increase during cranial neural fold elevation in the developing mouse midbrain

Formation of the mouse cranial neural tube requires the sequential elevation, apposition, and fusion of the cranial neural folds on either side of the midline (Figure 1A). The spatiotemporal forces that drive these changes, and how neuroepithelial integrity is maintained in the presence of these forces, are not well understood. To address these questions, we analyzed the spatiotemporal distribution of forces during neural fold elevation in the developing mouse midbrain. The active form of the myosin II regulatory light chain (phosphorylated MRLC or pMRLC) is enriched at interfaces parallel to the mediolateral axis during midbrain elevation, suggesting that forces are spatially regulated during closure (McGreevy et al., 2015; Brooks et al., 2020; Bogart and Brooks, 2025). We found that pMRLC planar polarity was present in late, but not early, stages of neural fold elevation, suggesting that these forces may also be temporally regulated (Figure 1B-1D, Figure 1—figure supplement 1A). To investigate this possibility, we performed laser ablation experiments to measure relative contractile forces in the lateral midbrain, using myosin IIB-GFP to target individual cell edges for ablation (Bao et al., 2007). The peak recoil velocity in response to laser ablation is estimated to be proportional to the tension acting on the edge immediately prior to ablation (Hutson et al., 2003; Farhadifar et al., 2007). We found that peak recoil velocities were significantly higher at edges aligned with the mediolateral axis than at edges aligned with the anterior-posterior axis (Figure 1E-L, see Supplementary File 1 for a summary of all n values and statistical analyses), consistent with the effects of larger, tissue-scale ablations (De La O et al., 2025). The peak recoil velocities at anterior-posterior and mediolateral junctions were both significantly higher in late elevation compared to similarly oriented edges in early elevation (Figure 1E-L). Treatment of embryos expressing the adherens junction marker GFP-Plekha7 (Shioi et al., 2017) with the Rho-kinase inhibitor Y-27632 to reduce actomyosin contractility eliminated the response to ablation, indicating that these forces require myosin II activity (Figure 1—figure supplement 1B-D). Together, these results demonstrate that mechanical forces at cell-cell junctions are planar polarized in the mouse cranial neural plate and significantly increase during neural fold elevation.

**Figure 1.**
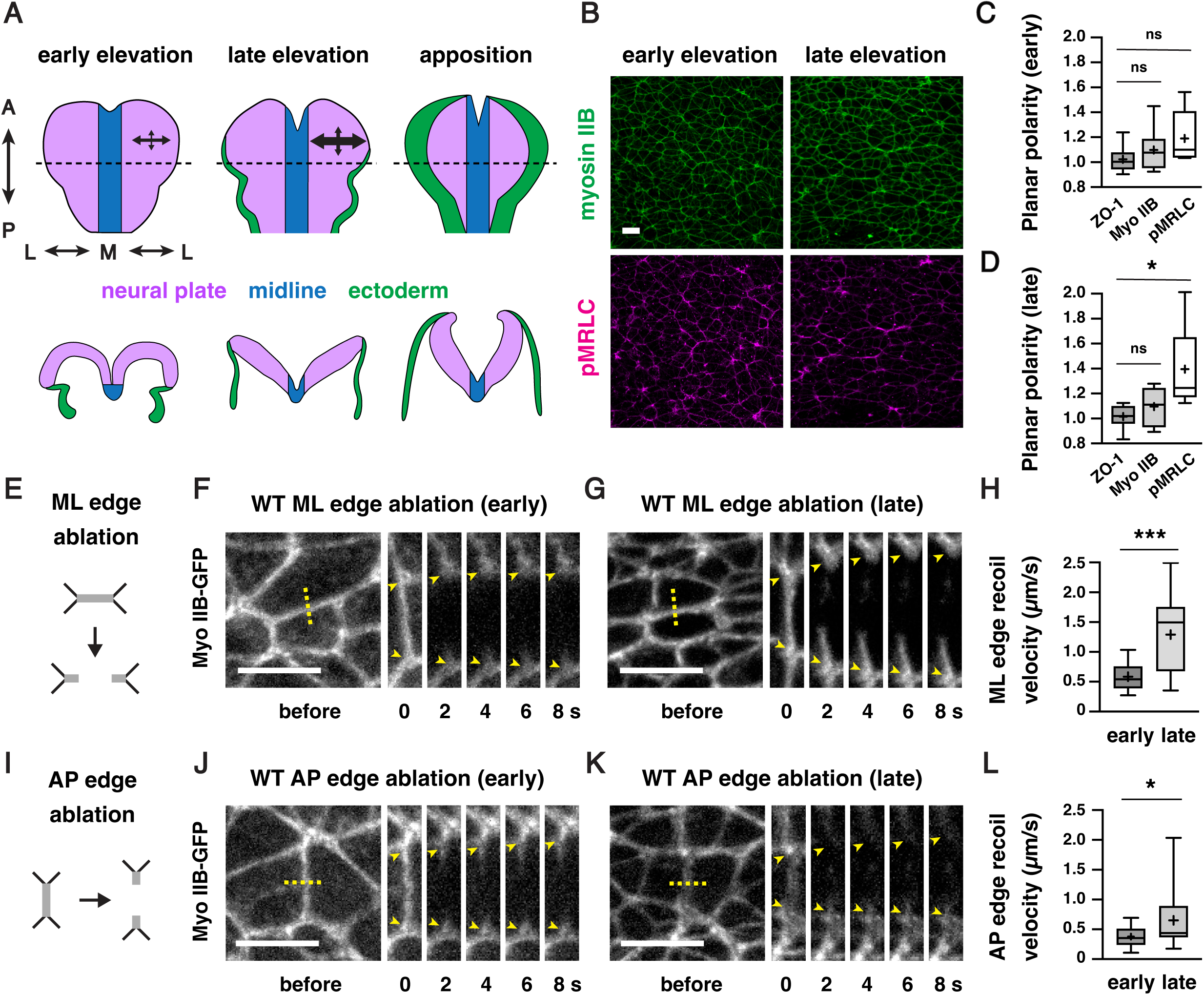
Actomyosin forces increase during cranial neural fold elevation. (A) Schematic of the presumptive midbrain and hindbrain of mouse embryos during early elevation (4-6 somites, E8.25), late elevation (7-9 somites, E8.5), and apposition (10-12 somites, E8.75). *En face* views (top), transverse views (bottom). Dotted lines, locations of transverse views. Arrows represent mechanical forces. (B) Localization of the myosin IIB heavy chain (Myo IIB) and the phosphorylated myosin II regulatory light chain (pMRLC) in early and late elevation. (C and D) pMRLC localization is planar polarized in late (D) but not early (C) elevation. Plots show mean fluorescence intensity at mediolateral (ML) edges (0-15° relative to the ML axis) divided by the mean intensity at anterior-posterior (AP) edges (75-90° relative to the ML axis). (E-G) Schematic (E) and kymographs (F and G) of ML edges before and 2-8 s after ablation in early (F) and late (G) elevation embryos expressing myosin IIB-GFP. (H) ML edge recoil velocity. (I-K) Schematic (I) and kymographs (J and K) of AP edges before and 2-8 s after ablation in early (J) and late (K) elevation embryos expressing myosin IIB-GFP. (L) AP edge recoil velocity. Boxes, 25^th^-75^th^ percentile; whiskers, 5^th^-95^th^ percentile; horizontal line, median; +, mean. 8 regions in 4 embryos in (C) and (D), 16-19 ablations in 6-9 embryos in (H and L). *p<0.04, ***p=0.0002 (Welch’s t-test). See Supplementary File 1 for for a summary of all data and statistical analyses. Maximum intensity projections, anterior up, edges oriented vertically in kymographs. Bars, 10 µm.

### Using embryonic stem cell-derived embryos to study neural tube closure

These results raised the question of how epithelial integrity is maintained as mechanical forces increase during midbrain neural fold elevation. We hypothesized that active mechanisms could be required to maintain cell and tissue structure in the presence of the forces required for cranial closure. Identifying key components of these processes requires direct live imaging of cell behavior in mutant embryos lacking critical force-response mechanisms. However, studying dynamic behaviors in mutant mouse embryos is challenging, as failure of cranial neural tube closure typically results in embryonic lethality, limiting the number of mutant embryos that can be obtained through genetic crossing strategies. To circumvent this limitation, we took advantage of established methods for generating mouse lines derived from mouse embryonic stem cells (ESCs) (Bradley et al., 1984; Beddington and Robertson, 1989; Nagy et al., 1990; Nagy et al., 1993; Hadjantonakis et al., 1998; Nagy and Rossant, 2001; Tam and Rossant, 2003). Injection of genetically modified mouse ESCs at the blastocyst stage (embryonic day 3.5, or E3.5) or aggregation with morula stage embryos are conventional methods for producing chimeric animals for generating stable mouse lines. By contrast, injection of mouse ESCs into host embryos at E2.5 (8-cell to morula stages) tends to produce embryos and animals in which nearly all epiblast lineages are derived from the injected ESCs, which can be used for immediate analysis (Lallemand and Brulet, 1990; Yagi et al., 1993; Poueymirou et al., 2007; Kiyonari et al., 2010; Ukai et al., 2017). Therefore, we sought to apply this system to generate embryos homozygous for lethal mutations to study the mechanisms of cranial neural tube closure.

To evaluate the reproducibility of this method for routine production of large numbers of genetically modified embryos with strong contributions from the injected ESCs, we performed a series of control experiments. First, GFP-negative parental ESCs (HK3i) (Kiyonari et al., 2010) were injected into E2.5 host embryos expressing Histone-H2B-GFP from the *ROSA26 (R26)* locus (Abe et al., 2011). Injected embryos were transferred to surrogate females and recovered for analysis at E8.5-9.5 to assess the contribution of the injected ESCs to the developing embryo during cranial closure (Figure 2A). As expected, the majority of embryos obtained with this method (90%) were derived nearly entirely from the GFP-negative ESCs in the embryonic lineages (Figure 2B). Host-derived cells expressing Histone-H2B-GFP were primarily observed in the gut tube, which retains a small number of extraembryonic cells at this stage (Kwon et al., 2008). These results confirm that mouse ESC injections provide an efficient strategy for generating embryos in which the embryonic lineages are nearly fully derived from the injected ESCs.

**Figure 2.**
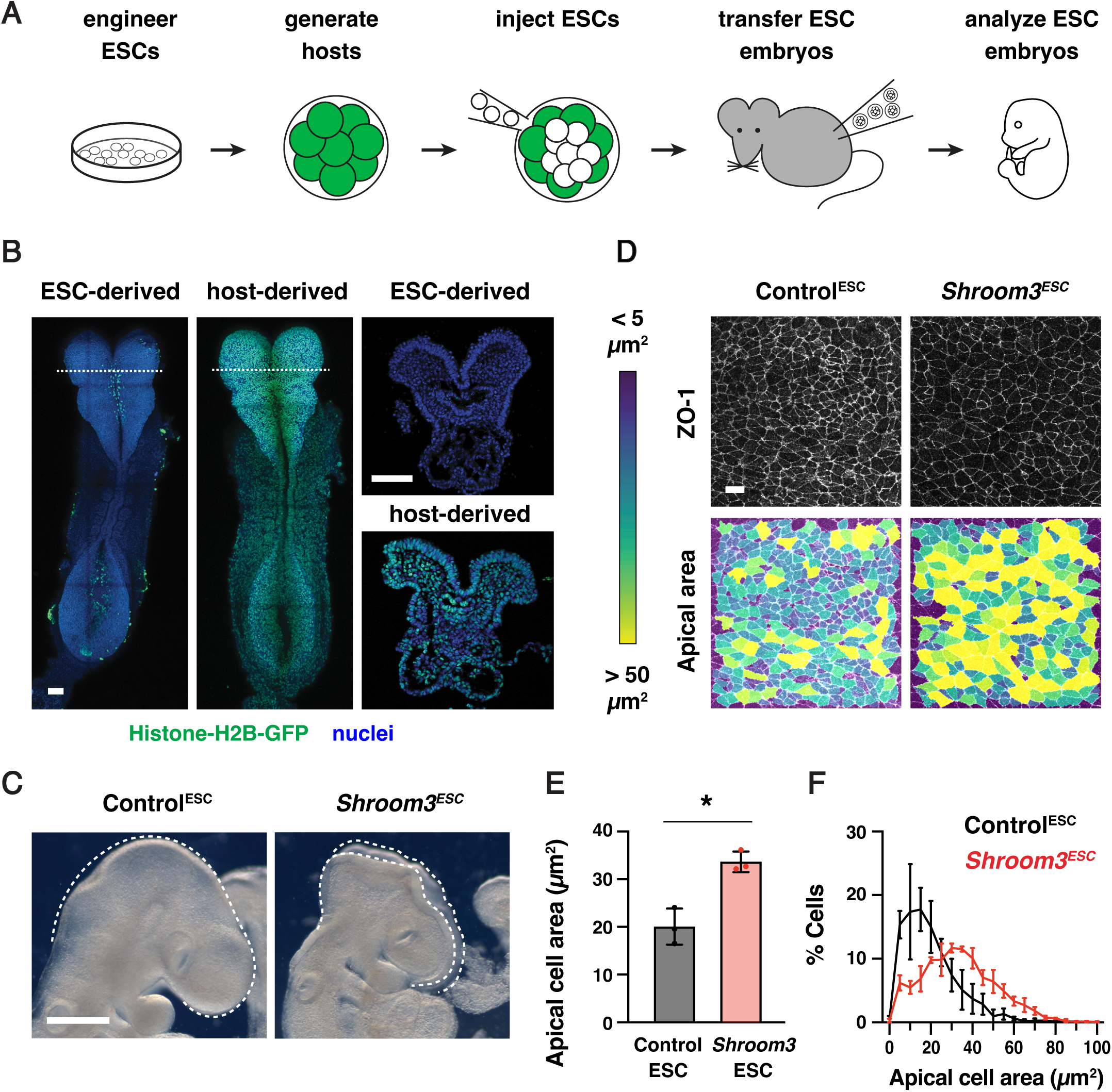
ESC-derived embryos recapitulate neural tube closure defects of *Shroom3* mutants. (A) Schematic of embryo generation from mouse ESCs. ESCs engineered using CRISPR/Cas9 gene editing were injected into E2.5 host embryos, ESC-injected embryos were transferred to surrogate females, and developing embryos were recovered for analysis. (B) A majority of embryos generated by ESC injection were derived primarily from GFP-negative ESCs (259/288 embryos), whereas some contained a significant contribution of Histone-H2B-GFP-positive host cells (29/288 embryos) and were excluded from further analysis. (C) Light micrographs of E9.5 Control^ESC^ and *Shroom3^ESC^* embryos (0/41 Control^ESC^ embryos and 79/79 *Shroom3^ESC^* embryos displayed exencephaly). Lateral views, dotted lines indicate the lateral edges of the cranial neural plate. (D) Lateral midbrain cells stained for ZO-1 (top) and color-coded by apical cell area (bottom). (E, F) Average apical cell area (E) and apical cell area distributions (F) of lateral midbrain cells in Control^ESC^ and *Shroom3^ESC^* embryos in mid-elevation (5-7 somites). Control^ESC^ embryos were derived from unedited (HK3i) or *Tyr* mutant ESCs. A single value was obtained for each embryo and the mean±SEM between embryos is shown (1572-2744 cells in 3 embryos/genotype). *p<0.02 (Welch’s t-test). Maximum intensity projections, anterior up. Bars, 100 µm (B), 500 µm (C), 10 µm (D).

To validate the use of this method for studies of cranial neural tube closure, we tested whether ESC-derived embryos reproduce the phenotypes of known closure mutants. Using CRISPR/Cas9 genome editing (Mulas et al., 2019), we generated ESCs homozygous for a null mutation in *Shroom3* (Figure 2—figure supplement 1A-C, Materials and methods), which encodes an actin-binding protein that is essential for cranial neural tube closure in mouse, frog, and chick embryos (Hildebrand and Soriano, 1999; Haigo et al., 2003; Nishimura and Takeichi, 2008; Massarwa and Niswander, 2013; McGreevy et al., 2015). The use of ESC injections to generate *Shroom3^ESC^*embryos produced an average of 6 *Shroom3^ESC^* embryos per surrogate female, a significantly increased yield of mutant embryos compared to heterozygote crosses. Embryos generated from unedited parental ESCs or from ESCs mutant for the *tyrosinase* coat color gene, referred to collectively as Control^ESC^ embryos, completed cranial closure normally, indicating that the parental ESC line and the ESC embryo generation protocol do not interfere with wild-type closure (Figure 2C). By contrast, *Shroom3^ESC^*embryos displayed a complete failure of cranial closure, resulting in fully penetrant exencephaly (79/79 *Shroom3^ESC^* embryos compared to 0/41 Control^ESC^ embryos). In addition, *Shroom3^ESC^* embryos showed significantly reduced apical constriction (Figure 2D-F), a Shroom3-dependent cell behavior that is essential for neural fold elevation (Haigo et al., 2003; McGreevy et al., 2015; Brooks et al., 2020; Baldwin et al., 2022). These results demonstrate that ESC-derived embryos recapitulate the cell behaviors and tissue-level closure defects characteristic of *Shroom3* mutants.

Next, we tested if this ESC injection approach could be used to generate embryos for live imaging. To visualize cell junctions, we derived ESCs from *R26-PHA7-EGFP* embryos that constitutively express the adherens junction marker GFP-Plekha7 from the *R26* locus (Shioi et al., 2017) and the established ESCs were injected into GFP-negative host embryos (Materials and methods). Imaging of the recovered embryos revealed widespread expression of GFP-Plekha7 throughout the cranial neural plate and confirmed that GFP-Plekha7 displayed the expected colocalization with N-cadherin (Figure 2—figure supplement 1D and E). A subset of embryos went on to produced viable and fertile adults that displayed germline transmission (Figure 2—figure supplement 1F). Together, these results demonstrate the utility of this method for generating numerically robust cohorts of genetically modified embryos for studying cellular and molecular mechanisms of cranial neural tube closure.

### Vinculin is required for cranial neural fold elevation

To determine how epithelial cells maintain cell adhesion and tissue integrity during cranial closure, we used this ESC-derived embryo method to generate embryos lacking the actin-binding protein Vinculin. Vinculin is recruited by mechanical forces to a variety of structures in cells, including adherens junctions, tight junctions, and cell-matrix adhesions (Goldmann, 2016; Bays and DeMali, 2017; Citi, 2019). Disruption of Vinculin expression or actin-binding activity leads to impaired cranial closure in the mouse embryo (Xu et al., 1998; Marg et al., 2010); however, how Vinculin influences neural tube closure is unknown. To elucidate the role of Vinculin in cranial closure, we used CRISPR/Cas9 gene editing to generate ESCs lacking Vinculin (Figure 3—figure supplement 1A and B). Homozygous *Vinculin* mutant ESCs were injected into 8-cell stage host embryos to generate *Vinculin^ESC^* embryos that lack Vinculin in the embryonic lineages, producing an average of 8 mutant embryos per surrogate female. In addition, ESCs heterozygous for a similar *Vinculin* deletion were injected to produce functionally wild-type Control^ESC^ embryos that retain Vinculin protein (Figure 3—figure supplement 1C). To test if these ESC-derived embryos reproduce the phenotypes of *Vinculin* mutants, we compared *Vinculin^ESC^* embryos to conditional mutants lacking *Vinculin* in the embryonic lineages generated by combining a conditional *Vinculin^flox^*allele (Zemljic-Harpf et al., 2007) with an epiblast-specific Sox2-Cre driver (Hayashi et al., 2002), hereafter referred to as *Vinculin^ΔEpi^*embryos. *Vinculin^ESC^* and *Vinculin^ΔEpi^* embryos displayed fully penetrant exencephaly at embryonic day E9.5, were slightly smaller than Control^ESC^ or littermate controls, and exhibited fully penetrant lethality by E10.5 (Figure 3—figure supplement 1D and E), recapitulating several known phenotypes of *Vinculin* mutants (Xu et al., 1998; Marg et al., 2010). Therefore, we used both *Vinculin^ESC^* and *Vinculin^ΔEpi^* embryos to study the effects of disrupting Vinculin activity in embryonic lineages on cranial neural tube closure.

To determine how Vinculin influences neural tube closure, we first examined when cranial neural plate defects arise in *Vinculin^ESC^* embryos. The midbrain neural folds elevate between E8.0 and E8.5, resulting in a decrease in the ratio of the apical span to the basal span of the tissue (Figure 3A-C). Neural fold elevation initiated normally in *Vinculin*^ESC^ embryos, but the neural folds failed to progress to late elevation and apposition, resulting in an arrest of neural tube closure (Figure 3A-C). The cell behaviors that drive neural tube closure require apical-basal polarity, which orients apical constriction and apical-basal elongation behaviors that are required for neural fold elevation (Grego-Bessa et al., 2015; McGreevy et al., 2015; Grego-Bessa et al., 2016; Brooks et al., 2020; Baldwin et al., 2022). F-actin enrichment at the apical cell contacts and basal localization of the extracellular matrix protein laminin occurred normally in *Vinculin^ESC^*embryos, indicating that Vinculin is not required to establish apical-basal polarity (Figure 3A). Moreover, although apical-basal elongation was slightly delayed in *Vinculin^ESC^* embryos, the final cell heights were comparable to Control^ESC^ embryos, demonstrating that Vinculin is also dispensable for apical-basal elongation (Figure 3—figure supplement 2A). The distribution of apical cell areas in *Vinculin^ESC^*embryos was indistinguishable from Control^ESC^ embryos in early elevation, indicating that apical constriction behaviors initiate normally (Figure 3D-G). By contrast, apical cell profiles in *Vinculin^ESC^*embryos were significantly larger in late elevation, suggesting a defect in the progression of apical constriction (Figure 3E and H). No differences in the frequency of mitosis or apoptosis were observed between *Vinculin^ESC^* and Control^ESC^ embryos, indicating that these defects are not due to a disruption of cell proliferation or survival (Figure 3—figure supplement 2B-E). Together, these results demonstrate that Vinculin is not required for cell proliferation, cell survival, apical-basal polarity, or apical-basal elongation in the midbrain neural plate, but Vinculin is required to maintain the apical constriction behaviors that are necessary for neural fold elevation.

**Figure 3.**
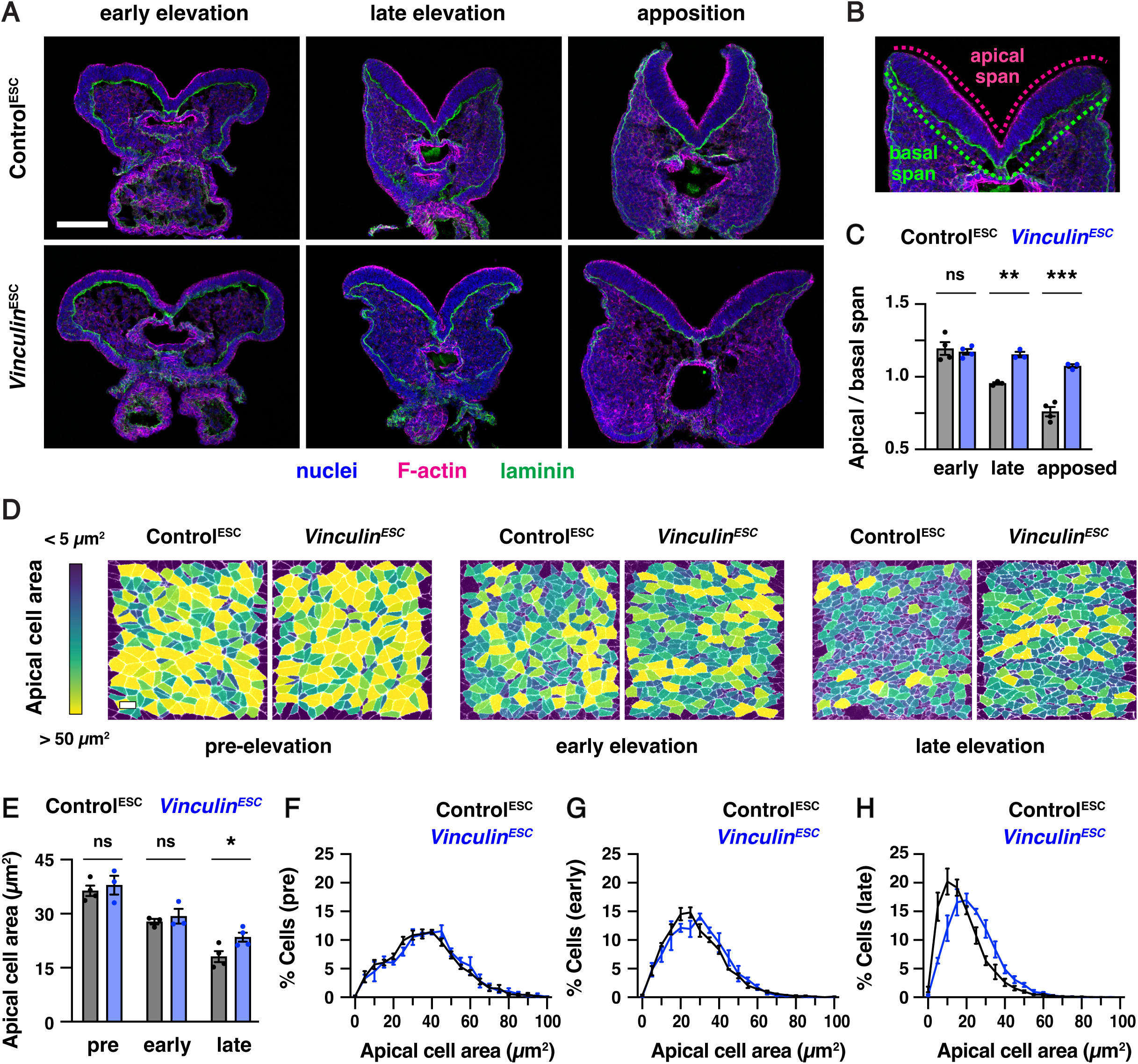
Vinculin is required for cranial neural fold elevation. (A) Transverse sections of the midbrain in Control^ESC^ and *Vinculin^ESC^* embryos in early elevation (4-6 somites), late elevation (7-9 somites), and apposition (10-12 somites). F-actin (phalloidin) labels the apical surface of the neuroepithelium and laminin labels the basal surface. (B) Schematic of apical and basal span measurements. (C) Apical-to-basal span ratios. (D) Lateral midbrain cells stained for ZO-1 (pre-elevation) or N-cadherin (early and late elevation) and color-coded by apical cell area. (E-H) Average apical cell areas (E) and apical area distributions (F-H) of lateral midbrain cells in Control^ESC^ and *Vinculin^ESC^* embryos before (F), early (G), and late (H) in elevation. A single value was obtained for each embryo and the mean±SEM between embryos is shown (1335-3955 cells in 3-4 embryos/genotype). *p<0.04, **p<0.004, ***p<0.001 (Welch’s t-test). Maximum intensity projections, apical up in A and B. Maximum intensity projections, anterior up in D. Bars, 100 µm (A), 10 µm (D).

### Vinculin is dispensable for force generation but is recruited to cell-cell junctions in a tension-dependent manner

As *Vinculin^ESC^*embryos display defects in neural tube closure as mechanical forces increase during neural fold elevation, we hypothesized that Vinculin could be required to generate or respond to the forces that drive neural tube closure. Consistent with a potential role in generating forces, Vinculin has been shown to regulate actomyosin localization at cell-cell junctions in epithelial cells in culture (le Duc et al., 2010; Leerberg et al., 2014) as well as in zebrafish cardiomyocytes and *Xenopus* embryos *in vivo* (Fukuda et al., 2019; van den Goor et al., 2024). Alternatively, Vinculin could stabilize cell adhesion in response to mechanical forces, as Vinculin is recruited to tricellular junctions that are predicted to be under increased tension (Cho et al., 2022; van den Goor et al., 2024) and acts to reinforce cell adhesion in the presence of experimentally applied forces *in vitro* and in the *Xenopus* embryo (Yonemura et al., 2010; Huveneers et al., 2012; Thomas et al., 2013; Kannan and Tang, 2015; Cho et al., 2022; van den Goor et al., 2024). However, whether Vinculin is required to generate or respond to endogenous forces during mouse neural tube closure is unknown.

To investigate whether Vinculin is required to generate mechanical forces during neural tube closure, we analyzed the localization of myosin IIB and filamentous actin (F-actin) in *Vinculin^ESC^* and *Vinculin^ΔEpi^* embryos. Myosin IIB and F-actin exhibit strong enrichment at cell-cell junctions in wild type embryos, including at bicellular junctions where two cells meet, tricellular junctions where three cells meet, and multicellular junctions where four or more cells meet (Figure 4A and B). By contrast, myosin IIB and F-actin were both present at cell-cell junctions in *Vinculin^ΔEpi^* embryos, but myosin IIB often appeared to dissociate from the membrane at tricellular and multicellular junctions, accompanied by a strong central accumulation of F-actin (Figure 4A and C). Similar results were observed in *Vinculin^ESC^* embryos (Figure 4— figure supplement 1A and 1B). Myosin IIB and F-actin localization was less strongly affected at bicellular junctions, although myosin IIB at these structures was more diffusely localized (Figure 4—figure supplement 1C and D). These results indicate that Vinculin is not required to recruit actomyosin networks to cell-cell junctions, but is necessary for their organization at these structures.

**Figure 4.**
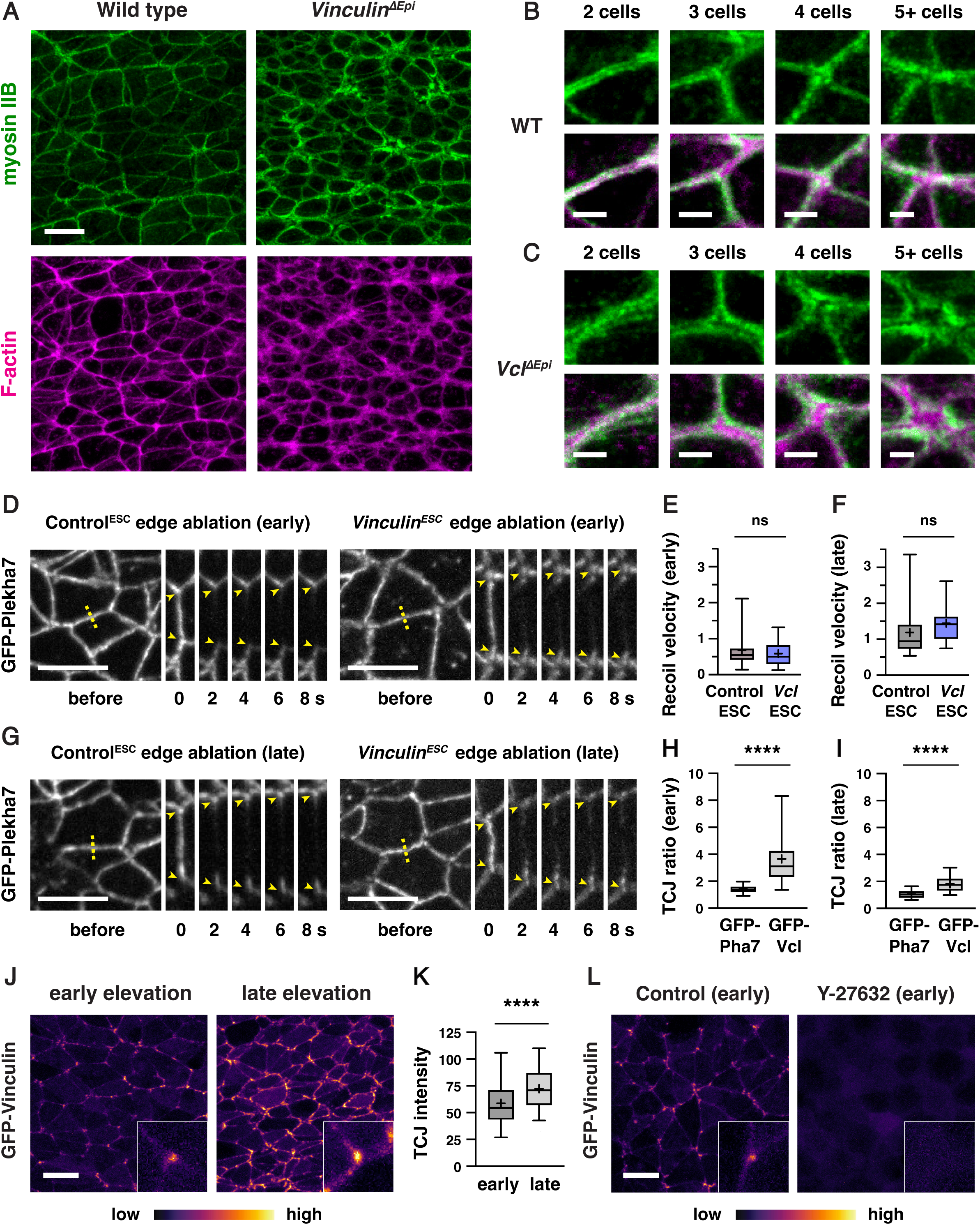
Vinculin is not necessary to generate force but is required for actomyosin organization at tricellular and multicellular junctions. (A) Localization of myosin IIB and F-actin (phalloidin) in late elevation wild-type and *Vinculin^ΔEpi^* embryos. (B, C) Close-ups of tricellular and multicellular junctions in wild-type (B) and *Vinculin^ΔEpi^*(C) embryos. (D-G) Laser ablation experiments in Control^ESC^ and *Vinculin^ESC^* embryos. (D, G) ML edges before and 2-8 s after ablation in early (D) and late (G) elevation embryos expressing GFP-Plekha7. (E, F) Peak recoil velocity after laser ablation of ML edges in early (E) and late (F) elevation. (H, I) GFP-Vinculin TCJ ratios (tricellular junction intensity divided by the mean intensity of the three connected bicellular junctions) in early (H) and late (I) elevation. (J, K) GFP-Vinculin localization in live wild-type embryos in early and late elevation. (L) Treatment of embryos for 2 h with 200 μm Rho-kinase inhibitor (Y-27632) in early elevation abolishes GFP-Vinculin localization at cell junctions. Boxes, 25^th^-75^th^ percentile; whiskers, 5^th^-95^th^ percentile; horizontal line, median; +, mean. 14-26 ablations in 5-7 embryos/genotype in E and F, 60-80 tricellular junctions in 3-4 embryos in H, I, and K. Boxes, 25^th^-75^th^ percentile; whiskers, 5^th^-95^th^ percentile; horizontal line, median; +, mean. ****p<0.0001, Welch’s t-test. Maximum intensity projections, anterior up, edges oriented vertically in kymographs. Bars, 10 µm (A, D, G, J, and L), 2 µm (B and C).

To test if Vinculin is required for the generation of actomyosin forces during neural fold elevation, we performed laser ablation of individual cell edges in wild-type and mutant embryos. As laser ablation requires live imaging of embryos before phenotypic differences between wild-type and mutant embryos become apparent, it can be challenging to obtain sufficient numbers of mutant embryos through traditional genetic crosses. Therefore, we took advantage of newly isolated ESCs expressing the adherens junction marker GFP-Plekha7 and used the same CRISPR/Cas9 strategy to generate a *Vinculin* null mutant ESC line expressing GFP-Plekha7 (Figure 4—figure supplement 2A). Mutant ESCs were injected into GFP-negative host embryos to generate *Vinculin^ESC^*embryos expressing GFP-Plekha7, which produced an average of 7 mutant embryos per surrogate female. The resulting embryos lacked Vinculin protein and displayed fully penetrant exencephaly, compared to Control^ESC^ embryos generated by injection of unedited GFP-Plekha7 cells, indicating that *Vinculin^ESC^* embryos expressing GFP-Plekha7 recapitulate the defects of *Vinculin* null mutants (Figure 4—figure supplement 2B and C). Moreover, analysis of relative forces using laser ablation revealed wild-type recoil velocities at mediolateral cell edges in *Vinculin^ESC^* embryos, with no differences compared to Control^ESC^ embryos in early or late elevation (Figure 4D-G). These results indicate that mechanical forces in the elevating neural plate are generated correctly in the absence of Vinculin.

As Vinculin is not required to generate forces in the cranial neural plate, we next asked if it is necessary to reinforce cell adhesion in response to mechanical forces during elevation. If Vinculin stabilizes cell adhesion under tension, then it is predicted to be recruited to high-tension structures within cells. To test this, we generated ESC-derived embryos that constitutively express N-terminally tagged GFP-Vinculin from the *R26* locus. GFP-Vinculin localized to cell-cell junctions in live-imaged embryos during early and late elevation and was enriched at tricellular junctions that are predicted to be under increased tension (Figure 4H-J). GFP-Vinculin intensity at tricellular junctions continued to increase during late elevation (Figure 4K), correlating with increasing mechanical forces, though its relative enrichment became less pronounced at these stages, likely due to increased recruitment to bicellular junctions. These changes were not due to changes in gene expression, as GFP-Vinculin levels did not change significantly during elevation (Figure 4— figure supplement 3). To test if GFP-Vinculin localization at tricellular junctions is regulated by force, we treated mouse embryos for 2 h with 200 μM of the Rho-kinase inhibitor Y-27632 to inhibit actomyosin contractility. Treatment with the Rho-kinase inhibitor abolished GFP-Vinculin localization at all junctions, indicating that Vinculin localization requires myosin activity (Figure 4L). These results demonstrate that Vinculin is recruited to cell-cell junctions in response to actomyosin contractility in the cranial neural plate, with the strongest localization observed at tricellular junctions that are under increased tension.

### Vinculin is required to maintain adherens junctions during elevation, but plays a minimal role in tight junction organization

The tension-dependent recruitment of Vinculin to cell-cell junctions, as well as the essential role of Vinculin in neural fold elevation, suggest that Vinculin may participate in functionally important force responses in the mouse cranial neural plate. Vinculin has well-known roles in stabilizing adherens junctions in the developing heart, skin, and vasculature (Zemljic-Harpf et al., 2007; Biswas et al., 2021; Kotini et al., 2022) and is also required for tight junction-mediated epithelial barrier function *in vitro* (Konishi et al., 2019) and in the *Xenopus* embryo (Higashi et al., 2016; van den Goor et al., 2024; Landino et al., 2025). However, whether Vinculin is required to regulate adherens junctions, tight junctions, or both during mouse neural tube closure is not known.

To investigate whether Vinculin regulates adherens junction localization in the cranial neural plate, we analyzed the localization of adherens junction proteins in Control^ESC^ and *Vinculin^ESC^*embryos (Figure 5A-D). The adherens junction protein N-cadherin was generally continuously localized along bicellular junctions, with few gaps in N-cadherin localization detected in either Control^ESC^ or *Vinculin^ESC^* embryos (Figure 5—figure supplement 1A and B). By contrast, gaps in N-cadherin localization were frequently observed at tricellular and multicellular junctions in *Vinculin^ESC^* embryos, particularly in late elevation when cells are under increased tension (35±4 gaps in *Vinculin^ESC^* and 18±5 gaps/region in Control^ESC^, mean±SD), whereas fewer defects in adherens junction localization were detected in early elevation when mechanical forces are lower (12±3 gaps/region in *Vinculin^ESC^*and 4±2 gaps/region in Control^ESC^) (Figure 5A-F, Figure 5— figure supplement 1C and D). Similar defects were observed in *Vinculin^ESC^* and *Vinculin^ΔEpi^*embryos, and mutant embryos generated with both methods displayed a loss of both N-cadherin and the adherens junction marker GFP-Plekha7 (Figure 6A and B, Figure 5—figure supplement 1E-G). To test if Vinculin is specifically required to regulate adherens junction localization under force, we analyzed protein localization at adherens junctions that are predicted to be under different levels of tension. Defects in GFP-Plekha7 localization were more frequent at higher-order junctions, with gaps detected at 18% of 3-cell junctions, 49% of 4-cell junctions, and 78% of 5+ cell junctions in *Vinculin^ESC^* embryos, compared to 5% of 3-cell junctions, 14% of 4-cell junctions, and 32% of 5+ cell junctions in Control^ESC^ embryos (Figure 5G-I). These results demonstrate that adherens junction defects increase during elevation and are most pronounced at junctions that sustain the highest forces, consistent with an essential role of Vinculin in maintaining cell adhesion under tension.

**Figure 5.**
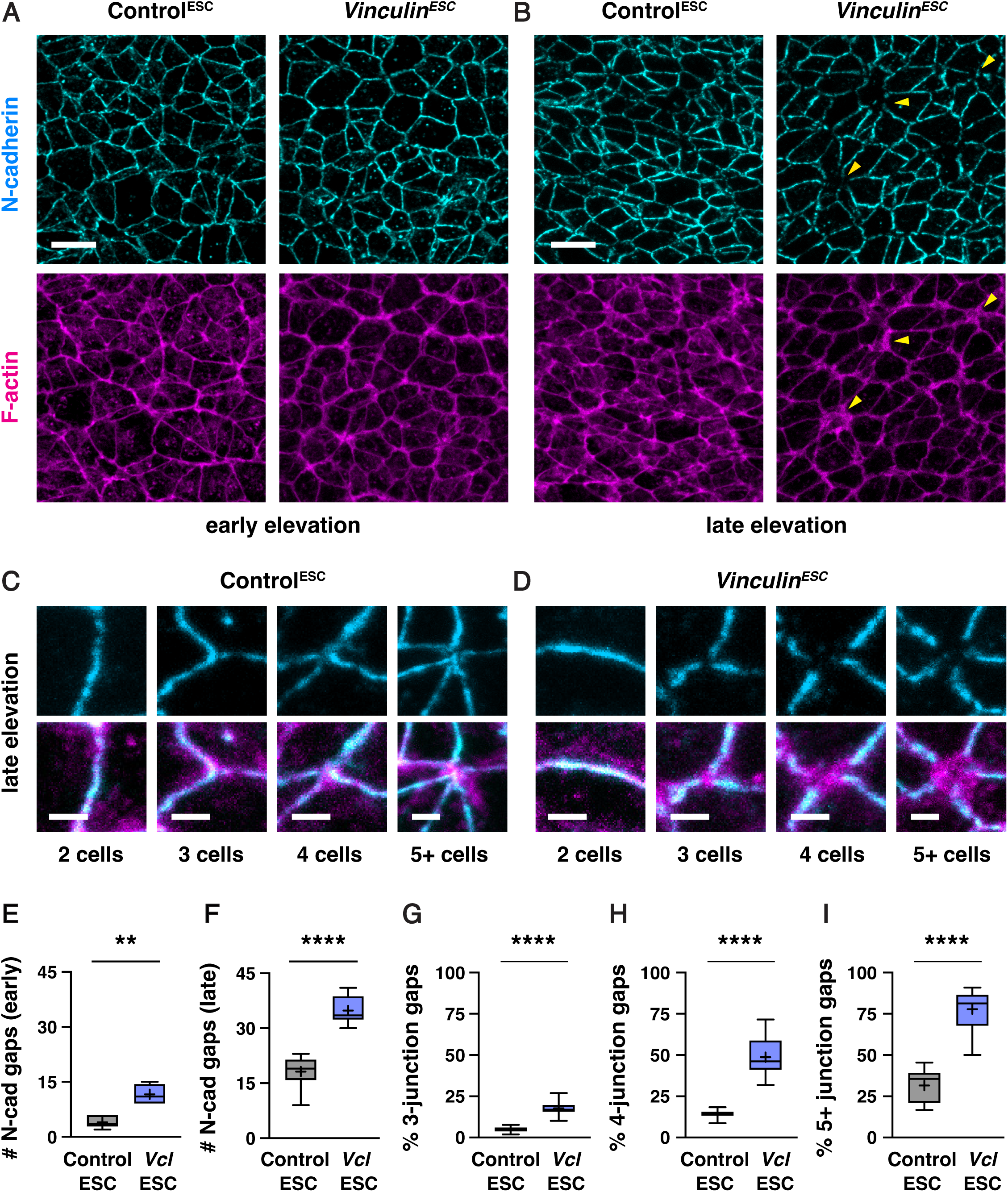
Vinculin is required for N-cadherin localization at tricellular and multicellular junctions. (A, B) Localization of N-cadherin and F-actin (phalloidin) in Control^ESC^ and *Vinculin^ESC^* embryos in early (A) and late elevation (B). Arrowheads indicate examples of multicellular junctions with gaps in N-cadherin localization. (C, D) Close-ups of bicellular, tricellular, and multicellular junctions in Control^ESC^ (C) and *Vinculin^ESC^* (D) embryos. (E, F) Number of gaps in N-cadherin localization in a 50 µm x 50 µm region of the lateral midbrain in early (E) and late (F) elevation. (G-I) Percentage of tricellular junctions (G), 4-cell junctions (H), and 5+-cell junctions (I) with gaps in GFP-Plekha7 localization. Boxes, 25^th^-75^th^ percentile; whiskers, 5^th^-95^th^ percentile; horizontal line, median; +, mean. 5-6 regions in 3 embryos in E and F, 81-175 3-cell junctions, 21-57 4-cell junctions, and 7-19 5+ cell junctions/region in 8-12 regions from 4-6 embryos in G-I. **p<0.002, ****p<0.0001 (Welch’s t-test). Maximum intensity projections, anterior up. Bars, 10 µm (A and B), 2 µm (C and D).

**Figure 6.**
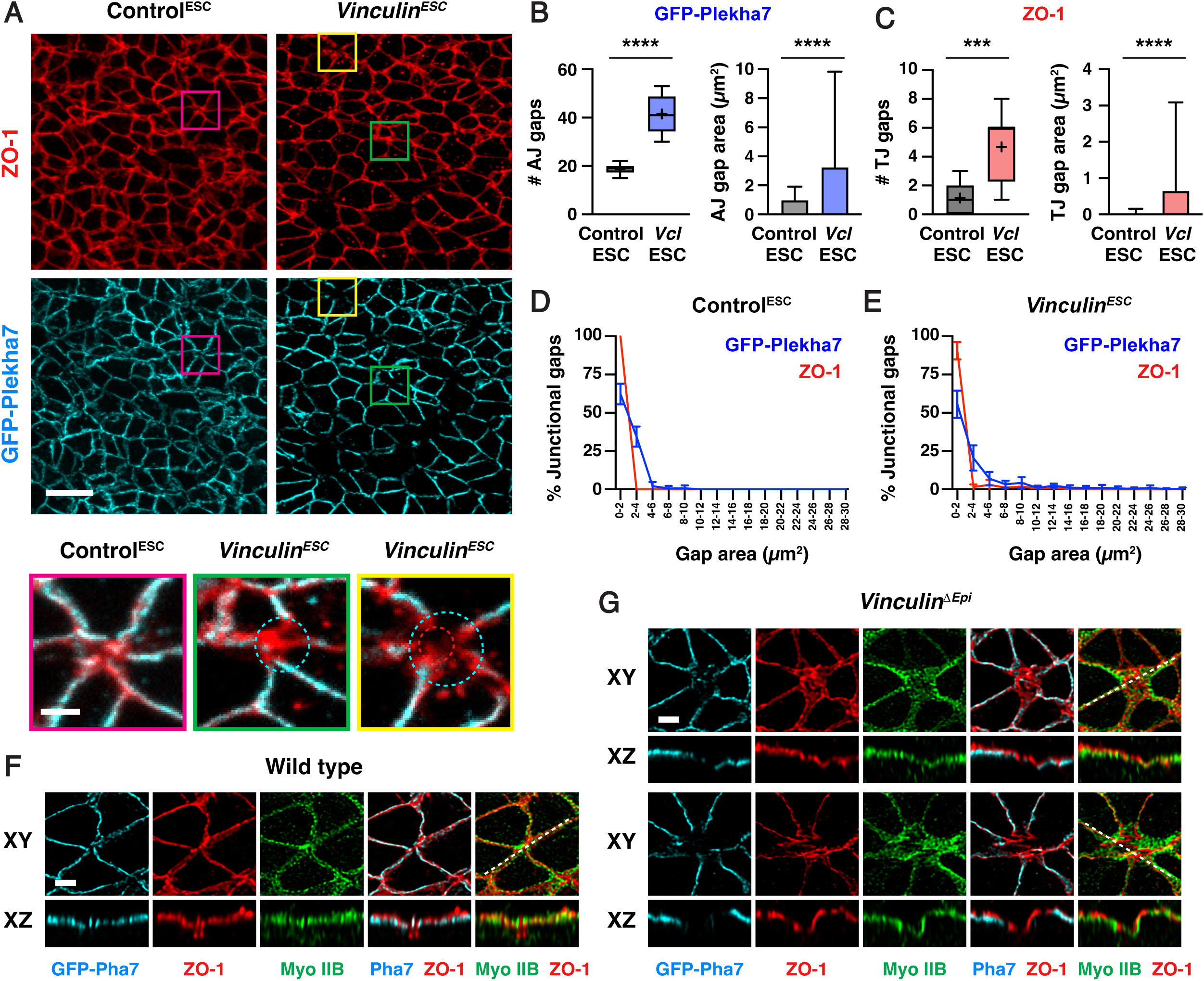
Loss of Vinculin disrupts tricellular and multicellular adherens junctions, whereas ZO-1 localization is generally maintained. (A) Localization of the tight junction protein ZO-1 and the adherens junction protein GFP-Plekha7 in late elevation Control^ESC^ and *Vinculin^ESC^* embryos. Bottom, merged images of the indicated regions showing a wild-type multicellular junction (magenta), a junction with a gap in GFP-Plekha7 localization (green), and a junction with a gap in GFP-Plekha7 and ZO-1 localization (yellow). Circles indicate gaps in GFP-Plekha7 (cyan) or ZO-1 (red) signal. (B, C) Number (left) and areas (right) of adherens junction gaps (B) and tight junction gaps (C) in a 50 µm x 50 µm region in late elevation Control^ESC^ and *Vinculin^ESC^* embryos. Note the differences in scale between the adherens junction and tight junction plots. (D, E) Area distributions of adherens junctions gaps detected with GFP-Plekha7 (blue) and tight junction gaps detected with ZO-1 (red) in Control^ESC^ (D) and *VInculin*^ESC^ (E) embryos. Five adherens junction gaps in (E) are outside of the x-axis range. (F, G) *En face* (XY) views and optically reconstructed (XZ) cross-sections of Airyscan z-stacks of GFP-Plekha7, ZO-1, and myosin IIB localization at multicellular junctions in late elevation wild type (F) and *Vinculin^ΔEpi^* (G) embryos. Boxes, 25^th^-75^th^ percentile; whiskers, 5^th^-95^th^ percentile; horizontal line, median; +, mean. Mean±SD between regions in B and C (right plots), D, and E. 150-498 adherens junction gaps in 8-12 regions from 4-6 embryos. ***p=0.0004, ****p<0.0001 (Welch’s t-test). Maximum intensity projections, anterior up in A, F, and G (top panels). Maximum intensity projections, apical up in F and G (bottom panels). Bars, 10 µm (A, top panels), 2 µm (A, bottom panels, F, and G).

As Vinculin is required to maintain adherens junction localization as mechanical forces increase during elevation, we next examined if Vinculin is required for the localization of tight junction proteins in the cranial neural plate. To investigate this possibility, we compared the distribution of GFP-Plekha7 with the tight junction-associated protein ZO-1. In contrast to adherens junctions, which displayed frequent gaps in GFP-Plekha7 localization in the absence of Vinculin, significantly fewer gaps were detected in ZO-1 localization (5±3 gaps in *Vinculin^ESC^* embryos and 1±1 gaps in Control^ESC^ embryos in late elevation) (Figure 6A-C, Figure 6—figure supplement 1A-C). These results indicate that tight junction gaps are less frequently detected than adherens junction gaps in both Control^ESC^ and *Vinculin^ESC^* embryos. Gaps in ZO-1 localization only occurred at sites where adherens junction localization were already defective, and were only present at a small subset of these sites (56/498 adherens junction gaps in *Vinculin^ESC^* embryos and 9/150 adherens junction gaps in Control^ESC^ embryos), and gaps in ZO-1 localization, when present, were consistently smaller than the gap in the corresponding adherens junction (Figure 6D and E). To better visualize these structures, we performed high-resolution Airyscan imaging and deconvolution of the apical junctional domain in Control^ESC^ and *Vinculin^ESC^* embryos. These results revealed that ZO-1 was continuously associated with the membrane at moderately sized gaps in adherens junction localization, whereas localized interruptions in ZO-1 signal were visible in regions where adherens junctions were more severely disrupted (Figure 6F and G). Despite these differences, ZO-1 signal remained slightly apical to GFP-Plekha7 at bicellular junctions in Control^ESC^ and *Vinculin^ESC^* embryos, indicating that these structures are correctly spatially delineated along the apical-basal axis (Figure 6F and G, Figure 6—figure supplement 1D and E). Together, these results indicate that adherens junctions are strongly disrupted in the absence of Vinculin, whereas tight junctions display only minor defects.

### Vinculin is required to maintain tissue integrity in response to force-intensive behaviors during neural tube closure

The findings that Vinculin is required to stabilize adherens junctions under tension, and that *Vinculin* mutants display fully penetrant defects in cranial neural tube closure, indicate that Vinculin regulates critical cell behaviors that drive structural changes in the mouse cranial neural plate. Multiple behaviors contribute to neural tube closure in mouse, chick, and *Xenopus* embryos, including apical constriction (Haigo et al., 2003; McGreevy et al., 2015; Chu et al., 2018; Brooks et al., 2020; Baldwin et al., 2022; Matsuda et al., 2023; Bogart and Brooks, 2025; Itoh et al., 2025), cell rearrangement (Davidson and Keller, 1999; Nishimura and Takeichi, 2008; Nishimura et al., 2012; Williams et al., 2014; Ossipova et al., 2015), and cell division (Morriss-Kay, 1981; Jacobson and Tam, 1982; Schoenwolf and Smith, 1990; Alvarez and Schoenwolf, 1991; Bogart and Brooks, 2025). However, how Vinculin influences cell behavior during cranial closure is unknown, in part due to the difficulty of directly visualizing dynamic cell behaviors in mutant embryos.

To address this question, we took advantage of the availability of Control^ESC^ and *Vinculin^ESC^* embryos expressing GFP-Plekha7 and used these embryos to perform time-lapse imaging of cell behavior in the lateral midbrain (Materials and methods). Control^ESC^ and *Vinculin^ESC^* embryos were mounted immediately before and during early elevation and imaged every six minutes for three hours (Videos 1-3). Control^ESC^ embryos displayed a 17±2% (mean±SEM) decrease in apical area after 1.5 hours and a 22±1% overall decrease in area after 3 hours (Figure 7A, Figure 7—figure supplement 1A and B). A similar change in area was observed after 1.5 hours in *Vinculin^ESC^* embryos, although junctional defects precluded an analysis of apical area at later time points in 3 of 5 mutant embryos (Figure 7A, Figure 7—figure supplement 1A and B). These results indicate that apical remodeling initiates correctly in the absence of Vinculin, but is often accompanied by defects in cell adhesion.

**Figure 7.**
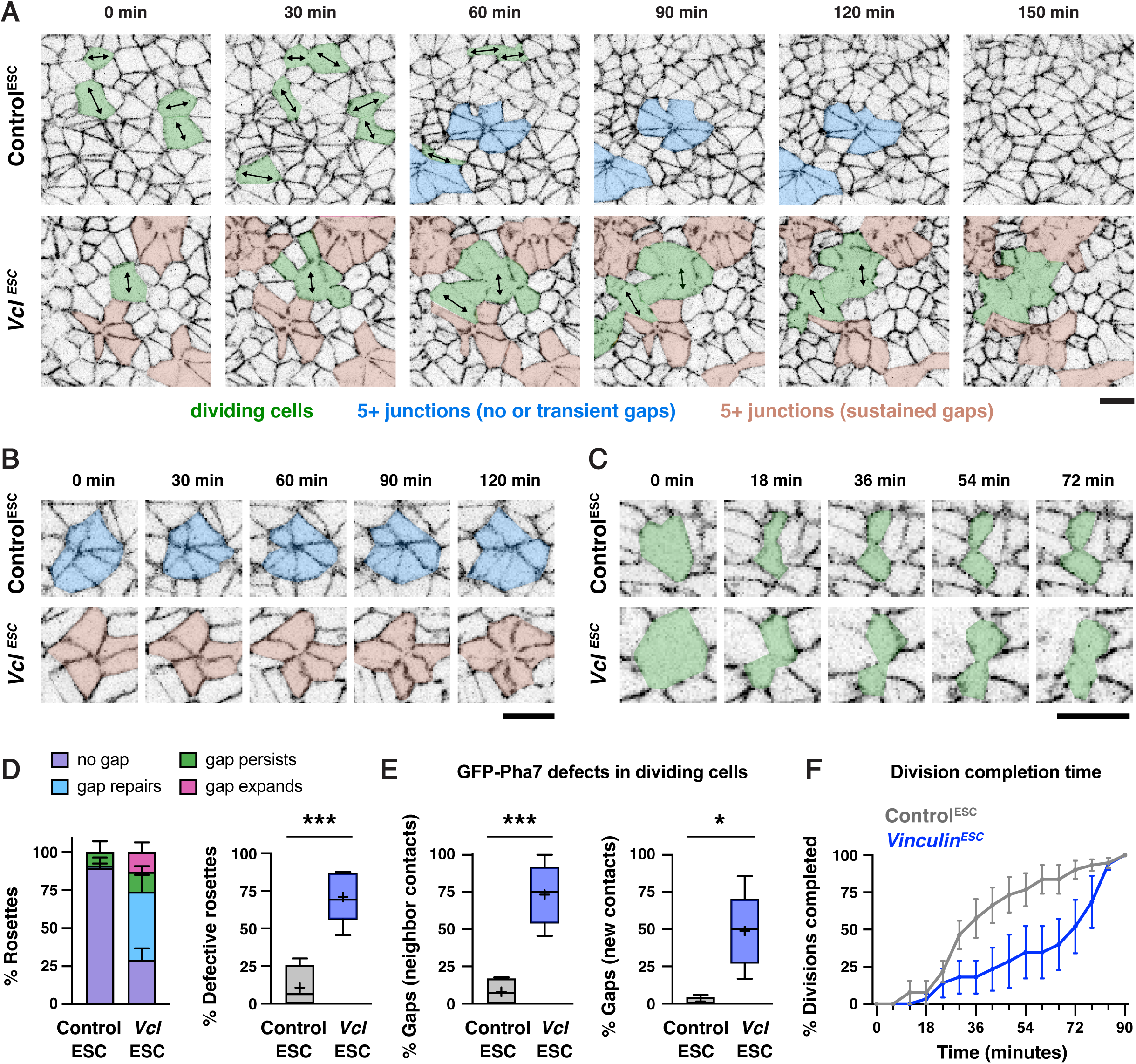
Vinculin is required to regulate force sensitive morphogenetic behaviors during elevation. (A) Stills from time-lapse movies of Control^ESC^ and *Vinculin^ESC^*embryos expressing GFP-Plekha7. Arrows, dividing cells. (B, C) Stills from time lapse movies showing rosettes (B) and dividing cells (C) in Control^ESC^ and *Vinculin^ESC^* embryos. t=0 in A and B is the start of the movie, t=0 in C is the time point immediately before the start of cleavage furrow ingression. (D) Left, percentage of rosettes with no gaps in GFP-Plekha7 signal or central gaps that repair, persist without changing size, or persist and increase in size in time-lapse movies of Control^ESC^ and *Vinculin^ESC^* embryos. Right, percentage of rosettes with gaps in Control^ESC^ and *Vinculin^ESC^* embryos. (E) Left, percentage of divisions with gaps in GFP-Plekha7 signal at interfaces between the dividing cell and neighboring cells that were not restored within 90 minutes. Right, percentage of divisions in which GFP-Plekha7 was not detected at a new vertex or interface within 90 minutes after the onset of cleavage furrow ingression. (F) Cumulative percentage of completed divisions that formed a new GFP-Plekha7-positive vertex or interface by the indicated times in Control^ESC^ and *Vinculin^ESC^* embryos. Embryos were imaged immediately before and during early elevation (2-5 somites). Boxes, 25^th^-75^th^ percentile; whiskers, 5^th^-95^th^ percentile; horizontal line, median; +, mean (in D right panel and E both panels). Mean±SEM between embryos in (D left panel and F). 47-48 rosettes in (D), 51-52 divisions in (E), and 28-50 divisions in (F) in 4-5 movies/genotype. *p<0.02, ***p<0.001 (Welch’s t-test). Maximum intensity projections, anterior up in A-C. Bars, 10 µm.

To investigate whether additional cell behaviors contribute to the junctional defects in mutant embryos, we analyzed higher-order junctions where 5 or more cells meet, also known as rosettes (Blankenship et al., 2006), which are thought to experience the highest forces in the tissue. Tracking of GFP-Plekha7 localization at these junctions revealed that 29/31 rosettes in Control^ESC^ embryos formed through cell rearrangement (the remaining 2 could not be determined), indicative of active remodeling. As expected based on our results in fixed embryos, few rosettes displayed central gaps in Control^ESC^ embryos (11±7%, mean±SEM, n= 5/47 rosettes), and the gaps that did form were either rapidly repaired (1/5 rosettes) or remained relatively small, with little or no further expansion in area during the imaging period (4/5 rosettes) (Figure 7A, B, and D). Rosettes were present in similar numbers in *Vinculin^ESC^* embryos, indicating that these structures form correctly in the absence of Vinculin (Figure 7—figure supplement 1C). However, the majority of rosette structures displayed gaps in GFP-Plekha7 signal (71±8%, n=33/48 rosettes) (Figure 7D), similar to the defects observed in fixed embryos (Figure 5I). Tracking rosettes over time in mutant embryos revealed that most gaps were either quickly repaired (20/33) or persisted but did not increase in size (7/33). However, a subset of gaps at rosette centers (6/33) expanded and merged with other gaps in the tissue (Figure 7A, B, and D). These results demonstrate that defects in GFP-Plekha7 localization in rosettes are more frequent in *Vinculin^ESC^* embryos, but are only occasionally associated with a widespread disruption of cell adhesion.

As the mislocalization of GFP-Plekha7 at higher-order junctions was only infrequently associated with a broader disruption of cell adhesion in the absence of Vinculin, we examined whether other cell behaviors could contribute to the defects in *Vinculin^ESC^* mutants. Cell division is known to exert mechanical forces on epithelial sheets through mitotic cell rounding (Kondo and Hayashi, 2013; Monster et al., 2021) and contraction of the cytokinetic ring (Founounou et al., 2013; Guillot and Lecuit, 2013; Herszterg et al., 2013). Vinculin is required to reinforce cell adhesion during these processes in MDCK cells and in the *Xenopus* embryo (Higashi et al., 2016; Monster et al., 2021; Landino et al., 2025). To test if Vinculin is required to maintain cell adhesion during cell division in the mouse cranial neural plate, we analyzed GFP-Plekha7 localization in dividing cells, which were identified by the presence of an ingressing cleavage furrow and tracked until the appearance of GFP-Plekha7 at the new vertex or interface (Figure 7A and C). Similar numbers of dividing cells were detected in Control^ESC^ and *Vinculin^ESC^* movies (Figure 7—figure supplement 1D), indicating that Vinculin is not required to initiate cell division. However, in contrast to cells in Control^ESC^embryos, which maintained continuous GFP-Plekha7 localization throughout division (Figure 7C and E), dividing cells in *Vinculin^ESC^*embryos frequently displayed reduced GFP-Plekha7 signal at contacts with neighboring cells (73±9% of divisions in *Vinculin^ESC^* embryos compared to 8±5% of divisions in Control^ESC^ embryos). In addition, nearly half of the cells that began ingression in *Vinculin^ESC^* embryos failed to establish a new GFP-Plekha7-positive vertex or interface within 90 minutes (49±11% of divisions compared to 1±1% of divisions in Control^ESC^ embryos) (Figure 7C and E) and those that did took significantly longer to establish adhesion (Figure 7F, Figure 7–figure supplement 1E). Strikingly, defects in dividing cells often appeared to be focal points that expanded to produce larger regions of disrupted GFP-Plekha7 signal. In particular, sites of GFP-Plekha7 mislocalization in dividing cells often merged with each other and with defective rosettes, resulting in widespread regions of junctional disruption in *Vinculin^ESC^*embryos (Figure 7A). Together, these live imaging studies demonstrate that Vinculin is required to maintain cell adhesion in the presence of multiple force-intensive cell behaviors during cranial neural tube closure.

## Discussion

Mechanical forces are essential for tissue remodeling, but how cells maintain cell adhesion in the presence of the dynamic forces that drive epithelial morphogenesis in mammals is not well understood. Here we use gene targeting and ESC-derived embryo generation to elucidate how cells respond to mechanical forces during mouse cranial neural tube closure. We show that actomyosin-mediated contractile forces increase during neural fold elevation, but Vinculin is not required for the generation of mechanical forces during this process. Instead, Vinculin is recruited to tricellular and multicellular junctions under stress and is required to maintain cell adhesion at high-tension sites as forces increase during neural tube closure. Live imaging of mutant embryos reveals that adhesion defects frequently initiate at high-order adherens junctions and in dividing cells, where mechanical forces are predicted to be highest, whereas tight junctions remain largely intact. Together, these results show that Vinculin is required to maintain cell adhesion during the force-intensive epithelial remodeling events that promote cranial neural tube closure, and demonstrate an essential role for Vinculin in transducing mechanical forces into the critical cell- and tissue-scale changes that shape mammalian tissue structure.

A wide range of molecular and biophysical functions have been ascribed to Vinculin based on studies *in vitro* and in cultured cells (Goldmann, 2016; Bays and DeMali, 2017; Citi, 2019), but the critical roles of Vinculin in cells exposed to physiological forces *in vivo* are just beginning to be elucidated. Our findings that Vinculin is required to maintain cell adhesion under tension in the mouse cranial neural plate are reminiscent of previous studies showing that Vinculin is required to maintain epithelial integrity in MDCK cells (Monster et al., 2021) and in the *Xenopus* gastrula (Higashi et al., 2016; van den Goor et al., 2023; Landino et al., 2025). However, Vinculin knockdown in the *Xenopus* embryo results in a depletion of F-actin and pMRLC at tricellular junctions (van den Goor et al., 2024), whereas Vinculin is dispensable for cortical actomyosin localization and force-generating activity in the mouse cranial neural plate. Moreover, Vinculin functions primarily at tricellular junctions in the *Xenopus* embryo (van den Goor et al., 2024), whereas loss of Vinculin in the mouse cranial neural plate causes more striking defects at multicellular junctions that are predicted be under increased force load. Additionally, Vinculin is required to maintain cell adhesion under endogenous forces in the mouse embryo, whereas ectopic forces are necessary to induce junctional defects in the *Xenopus* embryo (van den Goor et al., 2024), consistent with an increased mechanical demand on cells during neural fold elevation.

Notably, we observed frequent interruptions in adherens junction localization in the wild-type mouse neural plate, which affected over 10% of 4-cell junctions and nearly a third of rosettes during elevation. This suggests that cells continually repair transient gaps in adhesion that arise during the course of normal development. How cells achieve epithelial homeostasis in order to maintain tissue integrity during closure is opaque. In one model, Vinculin and other actin-junction linkers could reinforce adherens junctions under tension in the cranial neural plate, similar to the combinatorial roles of junction-actin linkers in maintaining cell adhesion under tension in the *Drosophila* embryo (Sawyer et al., 2011; Razzell et al., 2018; Rauskolb et al., 2019; Yu and Zallen, 2020; Perez-Vale et al., 2021; Sheppard et al., 2023). Alternatively, the presence of a continuous tight junction network could maintain epithelial integrity when adherens junctions are locally disrupted, facilitating ongoing junctional repair. Although tight junctions do not typically support strong adhesion (Citi, 2019), they have been shown to promote mechanical force responses that could actively maintain epithelial integrity (Spadaro et al., 2017; Otani et al., 2019; Cho et al., 2022; Nguyen et al., 2024). The mouse cranial neural plate may be a uniquely high-tension setting in which multiple force-intensive behaviors—including apical constriction, cell division, and junctional remodeling—are coordinately regulated in time and space across large populations of cells. The requirement for active mechanisms to reinforce cell adhesion as mechanical forces escalate during cranial closure may be among the reasons why cranial neural fold elevation is a frequent point of failure in mouse neural tube mutants (Harris and Juriloff, 2018).

Mouse ESCs have long been used as a powerful strategy for creating genetically modified mouse lines (Capecchi, 2005). The ability to generate embryos and animals that are nearly completely derived from ESCs unlocks the potential to perform immediate analysis of mutant phenotypes (Poueymirou et al., 2007; Lai et al., 2015), generate fluorescent embryos and animals for live imaging (Hadjantonakis et al., 1998; Wang et al., 2016), and create postnatal models of complex human diseases such as cancer (Dow and Lowe, 2012; Alonso-Curbelo et al., 2021; Burdziak et al., 2023). Here we use this method to bypass the constraints of Mendelian genetics in order to generate large numbers of lethal mutant embryos for phenotypic analysis. ESC-derived embryos complete embryogenesis and develop into viable and fertile animals, demonstrating that this pipeline does not interfere with neural tube closure or other essential processes, providing an excellent platform for studying embryonic and post-embryonic development. This approach significantly accelerates the generation of desired genotypes, is well-suited to investigating dynamic cell behaviors by live imaging, and can be used to produce theoretically unlimited yields of mutant embryos. This approach has several advantages over standard genetic crossing strategies. Early ESC injections yield mutant embryos at a rate that is several times higher than conventional genetic crosses without the need for often difficult and time-consuming crossing strategies, significantly reducing the animal burden without sacrificing the monumental power of mouse genetics to elucidate the causes of, and innovate treatments for, human disease. Additionally, increasingly powerful genome editing technologies (Wang et al., 2013) can be harnessed to generate complex genotypes in mouse ESCs in culture followed by functional analyses *in vivo*, making it possible to rapidly test targeted mutations *in vivo* with a level of control previously limited to gastruloid and *in vitro* models. By comparison, fully *ex vivo* embryo-like models share this flexibility and go even further in reducing animal use (Amadei et al., 2022; Tarazi et al., 2022; Yilmaz et al., 2025; Jorgensen et al., 2025), but currently do not complete embryonic development, with few embryos surviving to neural tube closure stages. ESC-derived embryos can be used to rapidly characterize candidate genes, protein sequences, and disease variants in a physiological context, perform live imaging of mutants with genetically encoded fluorophores and biosensors, and disrupt multiple genes simultaneously to study combinatorial gene functions and model diseases with a multifactorial etiology, as is the case for many neural tube defects (Wilde et al., 2014; Juriloff and Harris, 2018; Lee and Gleeson, 2020). Combining the power of genome engineering with high-throughput *in vivo* studies of cell dynamics will accelerate exploration of the large and interconnected network of factors that modulate cell interactions during mammalian development.

## Materials and methods

### Mouse strains

The following published strains were used in this study: *Vcl^flox^*(*Vcl^tm1Ross^*) (Zemljic Harpf et al., 2007) (Jackson Laboratory stock #028451), Sox2-Cre (*Edil3^Tg(Sox2-cre)1Amc^*) (Hayashi et al., 2002) (Jackson Laboratory stock 008454), *R26-Plekha7-GFP* (*R26-PHA7-EGFP*) (Shioi et al., 2017) (RIKEN Accession # CDB0261K), and Myosin IIB-GFP (Bao et al., 2007). All stocks used for genetic crosses were maintained on an FVB/N background (Jackson Laboratory stock 001800). *R26-Histone2B-GFP (R26-H2B-EGFP)* (Abe et al., 2011) (RIKEN Accession # CDB0238K), C57BL/6J (Jackson Laboratory stock #000664), and C57BL/6N (Jackson Laboratory stock #005304 or Charles River stock #027) were used to generate host embryos for ESC injection. *Vcl^ΔEpi^* embryos were generated by crossing Sox2-Cre; *Vcl^Δ/+^* males to *Vcl^flox/flox^* females to generate Sox2-Cre; *Vcl*^Δ*/-*^ mutant embryos. Controls for *Vcl^ΔEpi^* embryos were stage-matched wild type (*Vcl^+/+^* or *Vcl^flox/+^*) or heterozygous (*Vcl*^Δ*/+*^ or *Vcl^-/+^*) littermate controls. Embryos were harvested at E7.75-E10.0 and timed matings were confirmed by the presence of a vaginal plug, with noon of the day of the plug considered E0.5. Embryos were staged by somite number, corresponding to E8.0 (pre-elevation, 0-3 somites), E8.25 (early elevation, 4-6 somites), E8.5 (late elevation, 7-8 somites), and E8.75 (apposition, 9+ somites). For *Vcl^ΔEpi^* embryos and littermate controls, embryos were genotyped by PCR using the following primers to test for the *Vcl^flox^*and *Vcl*^Δ^ alleles: 5’-CCTGCGCGGGATTACCTCATTGAC-3’ (Vcl flox Fwd), 5’- TTACGCCTAGCACTTGAA-3’ (Vcl Δ Fwd), 5’- TGCTCACCTGGCCCAAGATTCTTT-3’ (Vcl common rev). The following primers were used to test for Sox2-Cre: 5’-CTTGTGTAGAGTGATGGCTTGA-3’ (WT Sox2 Fwd), 5’- CCAGTGCAGTGAAGCAAATC-3’ (Sox2-Cre Fwd), 5’-TAGTGCCCCATTTTTGAAGG-3’ (Common Rev).

All animal experiments were conducted in accordance with the Guide for the Care and Use of Laboratory Animals of the National Institute of Health and approved Institutional Animal Care and Use Committee protocols (15-08-013 and 90-12-033) of Memorial Sloan Kettering Cancer Center.

### Mouse embryonic stem cell (ESC) culture

ESC modifications were performed on the HK3i mouse ESC line derived from a C57BL/6N background (Kiyonari et al., 2010) or wild-type Plekha7-GFP ESC clone A3 derived from *R26-PHA7-GFP* embryos in C56BL/6N (Shioi et al., 2017). ESCs were grown on 0.1% gelatin-coated tissue culture plates in Cellartis 3i mES/iPSC culture medium (Takara Bio) containing ERK/MEK inhibitor, GSK3β inhibitor, FGFR-TK inhibitor, and 1000 U/mL ESGRO recombinant mouse LIF protein (EMD Millipore), referred to as 3i/LIF medium. HK3i mouse ESCs (Kiyonari et al., 2010) were grown in a humidified incubator at 37 °C with 5% CO_2_ on mouse embryonic fibroblast (MEF) feeder cells or drug-resistant MEF feeder cells (DR-4) (Tucker et al., 1997) that were mitotically inactivated by 4,000 cGy X-ray irradiation or 10 ug/mL mitomycin C for 2-3 hr. Feeder cells were plated in medium containing 1x Knockout DMEM (Gibco), 5-15% FBS (GeminiBio and Millipore Sigma), 2 mM GlutaMAX Supplement (Gibco), and 1 mM MEM non-essential amino acids (Gibco). ESC clones or non-clonal, pooled cell populations for DNA analyses to assess the occurrence of knock-in insertions or indel mutations were grown without feeder cells in medium containing 1x KO DMEM (Gibco), 15% KO serum replacement (Gibco), 2 mM GlutaMAX Supplement (Gibco), 1 mM MEM non-essential amino acids (Gibco), 1000 U/mL ESGRO recombinant mouse LIF protein (EMD Millipore), 1 uM 2-mercaptoethanol (Gibco), 1 uM PD0325901 (amsbio), and 3 uM CHIR99021 (amsbio), referred to as KSR/2i medium, in a humidified incubator at 37°C with 5% CO_2_.

### Plasmid vectors for genome editing in ESCs

To select the gRNA sequences to generate *Shroom3* and *Vinculin* knockout alleles, the following design tools were used: IDT, CRISPOR (Concordet and Haeussler, 2018), GuideScan2 (Perez et al., 2017; Schmidt et al., 2025), and CHOPCHOP v3 (Labun et al., 2019). DNA sequence information and chromosomal coordinates were obtained from Ensembl (Cunningham et al., 2021; Martin et al., 2022) and when a genome could be selected, *Mus musculus* GRCm39/mm39 was used. To generate plasmids expressing sgRNAs for ESC modification, DNA inserts containing sequences corresponding to two crRNAs, each of which was flanked by the U6 promoter and gRNA scaffold sequences, were *in vitro* synthesized and subcloned into the *pUC57* vector (GenScript). The 2x tandem U6-sgRNA fragment was then cloned into the *pPuro-Dest-Cas9* plasmid, a derivative of *pX330.puro* (*pX330.puro* was a gift from Sandra Martha Gomes Dias, Addgene plasmid #110403). The following gRNA target sites were selected for an ∼2,710 bp deletion within exon 5 of *Shroom3*: 5’ target GTACTCGAAGATGCTCGAAC and 3’ target TTCGCCAGCGGACGCCTAGT. The gRNA target sites to induce an ∼536 bp deletion with breakpoints within introns 2 and 3 flanking *Vinculin* exon 3 were 5’ (intron 2) GGCCAAGTCCTGTTATAAAT and 3’ (intron 3) CCGTTAAAATTGCACTTTAG-3’.

To generate the *pR26-GFP-4xGSS-Vcl-201* targeting vector, the *pR26-GFP-4xGSS* backbone containing the N-terminal GFP and an *in vitro* synthesized cDNA fragment containing the coding sequence from the *Vcl-201* isoform preceded by a 4xGSS linker sequence (Genscript) was first subcloned into the *pUC57* vector (*pUC57-4xGSS-Vcl-201*). The *4xGSS-Vcl* insert was excised and cloned into the *pR26-GFP* targeting vector backbone containing a 1,083 bp 5’ homology arm, an adenovirus major late transcript splice acceptor (Friedrich and Soriano, 1991), EGFP, a poly-A signal from the bovine growth hormone gene, a human Ubiquitin C gene promoter-driven Neo-R cassette, and a 811 bp 3’ homology arm.

### Generation of *Shroom3* mutant, *Vinculin* mutant, and *R26-GFP-Vinculin* ESC lines

Genome editing was performed using methods adopted from published protocols (Ukai et al., 2017; Mulas et al., 2019) in the HK3i cell line derived from a C57BL/6N background (Kiyonari et al., 2010) or a wild-type GFP-Plekha7 ESC line (clone A3) derived in-house from *R26-PHA7-EGFP* embryos (Shioi et al., 2017) according to previously described methods (Kiyonari et al., 2010). *Shroom3* mutant ESC clones 15, 21, 33, and 38 carried biallelic deletions of exon 5 of the *Shroom3-203* isoform, producing a frameshift mutation in downstream exons (Figure 2—figure supplement 1B). Control embryos for *Shroom3^ESC^* were generated from wild-type HK3i ESCs (Kiyonari et al., 2010) or *tyrosinase (Tyr)* mutant ESCs (clones 32 and 41) derived from HK3i generated by CRISPR/Cas9 genome editing. *Vinculin^ESC^* embryos were generated from the homozygous mutant *Vinculin* ESC clone 58, which carried biallelic deletions of exon 3 of the *Vinculin-201* isoform, producing a frameshift mutation in downstream exons that alters the protein sequence after amino acid 79 of 1,066 (Figure 3—figure supplement 1B, Figure 4—figure supplement 2A). *Control^ESC^*embryos were generated from the heterozygous *Vinculin* ESC clone 15, which carried a monoallelic deletion. *Vinculin^ESC^* embryos expressing GFP-Plekha7 from the *R26* locus were generated from *Vinculin* biallelic mutant ESC clone 41 and control embryos were generated from wild-type *R26-PHA7-EGFP* ESC clone A3.

To generate the *Shroom3* and *Vinculin* ESC lines, HK3i or R26-GFP-Plekha7 cells were transfected using Lipofectamine™ LTX Reagent with PLUS™ Reagent (Invitrogen) with *pPuro-(Dest-sgRNA)-Cas9* constructs that express the Puromycin resistance gene, Cas9, and corresponding sgRNAs per the manufacturer’s instructions. Lipofected cells were plated at 3 x 10^4^ to 1 x 10^5^ cells onto 60 mm tissue culture-treated dishes coated with 0.1% gelatin pre-seeded with DR-4 MEF feeder cells. Lipofected cells were subjected to puromycin selection (2 mg/ml) (Gibco) for 24 hr starting 18 to 24 hr after lipofection. Following puromycin removal, cells were cultured for an additional 8-12 days in non-selective 3i/LIF medium. Individual colonies were isolated into 96-well plates, dissociated, and expanded to generate the original frozen stocks (passage 0, P0 generation) and for DNA isolation. DNA extraction was performed using the DNeasy Blood and Tissue Kit (Qiagen). Candidate biallelic and monoallelic deletion mutant clones at each locus were identified by PCR screening using the indicated primers. Selected clones were expanded to P2 to P5 generations for embryo production by injection. Fractions of individual clones were also cultured in non-adhesive, floating cell clump-forming conditions for elimination of feeder cells. DNA isolated from floating ESC clumps (DNeasy Blood and Tissue Kit) was PCR amplified for deep sequencing of amplicons around the deletion (for sequencing mutant alleles) or around the 5’ and 3’ CRISPR cut sites (for sequencing heterozygous controls), with 97,000-850,000 reads obtained/amplicon (Sloan Kettering Integrated Genomics Operation Facility) using the indicated primers.

Primers used to screen for *Shroom3* deletions were: A (forward) CCCTGCCATCTCCTTTCTCCTG, B (reverse) CCGTCTTGACGTAGCGGATGTC, G (forward) CCCTGCCATCTCCTTTCTCCTG, and D (reverse) CCCTGCCATCTCCTTTCTCCTG. Primer pair A and B amplifies across the 5’ gRNA target site and is predicted to generate an amplicon of ∼232 bp for the wild-type allele only. Primer pair G and D amplifies across the 3’ gRNA target site and is predicted to generate an amplicon of ∼246 bp for the wild-type allele only. Primer pair A and D amplifies across the deletion and is predicted to generate an amplicon of ∼269 bp for deletion alleles only.

Primers used to screen for *Vinculin* deletions were: A (forward) TCACAAGCCTAGTGCACAGAG, B (reverse) TCACAAGCCTAGTGCACAGAG, C (forward) GTGGGTGCCAAGAACCAAAC, and E (reverse) GTTTCACTGTGTAGCCTTGGC. Primer pair A and B amplifies across the 5’ gRNA target site and is predicted to generate an amplicon of ∼238 bp for the wild-type allele only. Primer pair C and E amplifies across the 3’ gRNA target site and is predicted to generate an amplicon of ∼236 bp for the wild-type allele only. Primer pair A and E amplifies across the deletion and is predicted to generate an amplicon of ∼241 bp for deletion alleles and ∼777 bp for the wild-type allele.

To generate the *R26-GFP-Vinculin* ESC knock-in line, HK3i ESCs were transfected using Lipofectamine™ LTX Reagent with PLUS™ Reagent (Invitrogen) with the *pDonor-MCS-ROSA26* vector (Addgene 37200) (Perez-Pinera et al., 2012) containing ∼0.8 kb left and right homology arms (chromosome 6: GRCm39:6:113052181:113053829) for targeted insertion of a GFP-Vinculin cDNA and a neomycin cassette flanked by FRT sites into the *R26* locus. The donor plasmid was transfected with a *pPuro-(Dest-sgRNA)- Cas9* vector (sgRNA R26-T2) (Nakao et al., 2016) to induce a double-strand break in the *R26* locus. Lipofected cells were plated at 1 x 10^5^ onto 60 mm tissue culture-treated dishes coated with 0.1% gelatin containing DR-4 MEF feeder cells. Cells were subjected to puromycin selection (2 mg/ml) (Gibco) for 24 hr starting 18-24 hr after lipofection. Following puromycin removal, cells were treated with 200 ug/uL Geneticin (G418) (Gibco) for 48 hr and this was decreased to 150 ug/uL G418 for the following 6-10 days. Individual colonies were isolated into 96-well plates, dissociated, and expanded to generate the original stocks and for DNA isolation. DNA extraction was performed using the DNeasy Blood and Tissue Kit (Qiagen) or the 96-well plate method according to a protocol from the McManus lab (https://mcmanuslab.ucsf.edu/protocol/dna-isolation-es-cells-96-well-plate) modified in-house for PCR tube strips. Targeted ESC clones were identified by PCR screening using the indicated primers and copy number analysis using droplet digital PCR (Thermo Scientific) was performed to assess potential multi-copy integration and random integration events. *R26-GFP-Vinculin* clone 27 was expanded to P2 to P4 generations and used for embryo production by injection.

Primers used to screen for *R26* insertions were: RosaP1 (forward) CTCAGAGAGCCTCGGCTAGGTAGG, EGFPSQ1 (reverse) AGCTCCTCGCCCTTGCTCACC, iNeoSQ1 (forward) AGGAACTTCGTTGGTACCGTACG, and RosaP4 (reverse) GGAGACATCCACCTGGAAACCATTAATGG. Primer pair RosaP1 and EGFPSQ1 amplifies across the 5’ knock-in insertion junction and is predicted to generate an amplicon of 1,442 bp for the knock-in allele. Primer pair iNeoSQ1 and RosaP4 amplifies across the 3’ knock-in insertion junction and is predicted to generate an amplicon of 903 bp for the knock-in allele.

### Generation of ESC-derived embryos

Embryos were generated from selected clones through ESC injection into E2.5 (8-cell/morula stage) host embryos, followed by same-day transfer into E0.5 pseudopregnant females or next-day transfer into E2.5 pseudopregnant females according to standard methods (Behringer et al., 2014). E7.75-E10 embryos were recovered for analysis. Homozygous R26-Histone-H2B-GFP (R26-H2B-EGFP) males (Abe et al., 2011) and C57BL/6 females were mated to produce host embryos for ESC injection (Behringer et al., 2014). GFP-negative ESCs were injected into R26-Histone-H2B-GFP host embryos (Abe et al., 2011) and embryos with no GFP-positive host cells detected outside of the gut tube were analyzed. GFP-positive ESCs were injected into E2.5 wild-type C57BL/6 host embryos and embryos with widespread GFP expression in the cranial neural plate were analyzed. B6CBAF1 female mice were used to generate pseudopregnant surrogates. The analysis of the contribution of Histone-H2B-GFP-expressing host cells in Figure 2B was performed in *Vinculin^ESC^* and Control^ESC^ embryos.

### Whole-mount immunofluorescence

Embryos were dissected in ice cold phosphate buffered saline (PBS) and fixed in 4% paraformaldehyde (4% PFA, Electron Microscopy Services) for 1-2 hr at room temperature or overnight at 4 °C or with Dent’s fixative (4:1 methanol:DMSO) and fixed overnight at 4 °C. Dent’s-fixed embryos were rehydrated for 30 minutes each in solutions of 75:25, 50:50, and 25:75 methanol:PBS at room temperature. After fixation, embryos were rinsed three times in PBS + 0.1% Triton-X100 (PBS-Tr) and washed 3 x 30 min in PBS-Tr at room temperature. Embryos were then incubated in blocking solution (PBS + 0.1% Triton-X100, 3% BSA) for 1 hr at room temperature and incubated overnight at 4 °C in primary antibody diluted in antibody staining solution (PBS + 0.1% Triton-X100, 1.5% BSA). Primary antibodies used for staining of paraformaldehyde-fixed embryos were chicken anti-GFP to visualize Histone-H2B-GFP (Abcam ab13970, 1:2000), rabbit anti-N-cadherin (Cell Signaling 13116, 1:500), rabbit anti-nonmuscle Myosin IIB heavy chain (BioLegend 909901, 1:500), mouse anti-phospho-histone H3 (Ser10) (Cell Signaling 95777, 1:500), rabbit anti-phosphomyosin light chain (Thr18/Ser19) (Cell Signaling 95777, 1:100), rabbit anti-Shroom3 (UPT132, 1:100) (Hildebrand, 2005), and mouse anti-ZO1 (Invitrogen 33-9100, 1:500). Th rat anti-ZO-1 primary antibody (DSHB R26.4C, 1:100) was used to stain embryos fixed with Dent’s fixative (Figure 2D-F). Embryos were washed 3 x 30 min in PBS-Tr, incubated with AlexaFluor-conjugated secondary antibodies (1:500), AlexaFluor-546 Phalloidin (Thermo Fisher, 1:500), and/or Hoechst 33342 (1 mg/mL, Invitrogen, 1:1000) in antibody solution for 90 min at room temperature, washed 3 x 30 min in PBS-Tr, and stored in PBS-Tr at 4 °C until imaging. Embryos were mounted for imaging in Attofluor cell chambers (ThermoFisher A7816) by slightly compressing the embryos under pieces of coverglass held in place by vacuum grease (Dow Corning) on a circular 25 mm, #1.5 coverslip (Fisher NC1272770).

### Cryosectioning

Fixed embryos were equilibrated in 15% sucrose in PBS, then 30% sucrose in PBS for 30 minutes each with gentle rocking, mounted in blocks of OCT (Tissue-Tek) with the cranial flexure up, then tilted at an angle of 25-40° with the forebrain facing up to obtain transverse sections of the posterior midbrain. OCT blocks were frozen and 12 µm sections were acquired on SuperFrost Plus slides (Fisher Scientific) with a Leica CM3050S cryostat and kept frozen at -80 °C until analysis. For staining, slides were defrosted and sections were rehydrated with PBS + 0.1% Tween-20 (Sigma) (PBS-Tw) for 10 min, permeabilized with 0.5% PBS-Tr for 30 mins, and washed 5 x 5 min in PBS-Tw. Sections were incubated in blocking solution (PBS-Tw with 2% BSA) for 10 min at room temperature and incubated in primary antibody diluted in blocking solution overnight at 4°C. Primary antibodies were chicken anti GFP to visualize Histone-H2B-GFP (Abcam ab13970, 1:2000) and rabbit anti-laminin (Sigma L9393, 1:1000). Sections were washed 5 x 10 min with PBS-Tw, re-blocked for an additional 10 min in blocking solution, and incubated with AlexaFluor-conjugated secondary antibodies (1:500) with AlexaFluor-546 Phalloidin (Thermo Fisher, 1:500) and Hoechst 33342 (1 mg/mL, Invitrogen, 1:1000) in blocking solution for 30 min at room temperature. Embryos were washed 3 x 5 min with PBS-Tw and mounted under a #1.5 coverslip (Corning 2980-245) in fluorescence mounting media (Dako).

### Live embryo culture

All steps were performed in media equilibrated to 37 °C and 5% CO_2_. Timed pregnant females were euthanized and uterine horns were dissected and transferred into DMEM/F-12, GlutaMAX (DMEM) (ThermoFisher). Intact egg cylinders were removed from the decidua and transferred to 1:1 DMEM and whole embryo culture rat serum (Envigo). The parietal yolk sac and Reichert’s membrane were carefully dissected, maintaining an intact ectoplacental cone. The yolk sac and amnion were removed from around the neural folds, being careful to avoid damage to embryonic tissue. For live imaging, embryos were dissected immediately before and during early elevation (2-5 somites), mounted with the midbrain and hindbrain neural folds facing down in a 35 mm Lumox culture dish (Sarstedt) and immobilized with cut pieces of 70 µm cell strainer (Falcon) held in place with vacuum grease. Mineral oil (Sigma) was added to culture media to prevent evaporation. Embryos were incubated in a stage-top incubator (Pecon) at 37°C and 5% CO_2_ while imaging. For Rho-kinase inhibitor treatments, intact egg cylinders were treated with 200 µm Y-27632 (Sigma) or an equal volume of ultrapure H_2_O, pre-warmed in 1:1 DMEM:rat serum, and cultured at 37 °C and 5% CO_2_ in a 24-well Lumox culture plate (Sarstedt) for 2 hr before laser ablation or imaging.

### Microscopy

Cryosections were imaged on a Zeiss LSM700 or Zeiss LSM900 confocal microscope using a 20x/0.8 Plan-Apochromat objective with a 0.5x optical zoom and z-stacks of 8-14 µm were obtained with 2 µm z-slices and 1 µm z-steps. Whole mount fixed imaging was performed on a Zeiss LSM900 using a 20x/0.8 Plan-Apochromat objective or a Plan-Apochromat 40x/1.3 oil immersion objective. Z-stacks of 50-210 µm were obtained with 1.6-2.0 µm z-slices and 0.8-1.0 µm z-steps with an optical zoom of 0.45-0.5x for the 20x objective. Z-stacks of 10-100 µm were obtained with 0.3-0.5 µm z-slices and 0.6-1.0 µm z-steps with an optical zoom of 0.5-2.0x for the 40x objective. Tiled images were computationally stitched with 10% overlap using Zen-Blue software (Zeiss). Airyscan imaging (Wu and Hammer, 2021) was performed on a Zeiss LSM900 confocal microscope equipped with an Airyscan2 detector using a Plan-Apochromat 63x/1.4 oil immersion objective with an optical zoom of 1.7x. Z-stacks of 5-8 µm were obtained with 0.13 µm z-steps. Airyscan images were deconvolved using Huygens Professional Software (Scientific Volume Imaging). Live imaging was performed on a Zeiss LSM900 using a Plan-Apochromat 40x/1.3 oil immersion objective with an optical zoom of 0.5X. Z-stacks of 25-55 µm were obtained with 0.3-0.8 µm z-slices and 0.6-0.8 µm z-steps. Images were acquired at 6 min intervals for 3 hours. Light micrographs were acquired on a Zeiss Stemi 508 stereomicroscope with an attached Canon EOS T7i camera.

### Laser ablations

Laser ablation experiments were performed on a spinning disk confocal with a Yokogawa CSU X1 scan head and an Excelitas PCO.Edge 4.2 sCMOS camera on a Zeiss Observer Z1 microscope with a Zeiss 40X Plan NeoFluor 1.3-NA objective using an iLas ablation system (Gataca Systems, Massy, France). Images of the lateral midbrain were acquired as z-stacks of five z-slices with a step size of 0.5 μm using a Zeiss 40X Plan NeoFluor 1.3-NA oil-immersion objective and a PCO Edge 4.2 camera. To ablate a cell interface, 10 pulses of 355-nm light were focused at a single cell interface visualized with myosin IIB-GFP or GFP-Plekha7, along a 16-pixel line perpendicular to the cell interface. Pre-and post-ablation images were acquired every 2 s. The distance between the two tricellular junctions attached to the cut interface was measured immediately before and up to 8 s after ablation and the instantaneous velocity was measured at every time point. The peak velocity, which is predicted to correlate with tension at the junction prior to ablation, was used for analysis. Calibration of the iLas ablation system was performed prior to every experiment. Ablations that damaged edges other than the targeted edge or showed no measurable recoil were discarded. A maximum of 5 ablations were performed/neural fold and ablations were performed on both neural folds.

### Western blots

Single embryos staged between E8.0-9.5 were lysed for 10 min in ice cold RIPA buffer with Protease/Phosphatase Inhibitor cocktail (Cell Signaling) and manually homogenized with a pestle. Protein concentration was determined by BCA (Thermo Fisher). 30 µg of protein was boiled for 10 min in SDS sample buffer, separated on a PVDF membrane (Millipore), blocked in 5% milk in TBST, and immunoblotted with mouse anti-vinculin (Sigma, V9131, 1:1000) or mouse anti-β-catenin (BD Bioscience, 610153, 1:2000) antibodies overnight at 4 °C in 5% BSA, TBS-Tw. Membranes were washed 6 times for 5 mins with TBS-Tw, blotted with HRP-conjugated secondary antibodies (Jackson Laboratory, 1:5000) in 5% milk, TBST for 30 min at room temperature, and developed with Amersham ECL Western Blot Detection Reagent (Fisher Scientific).

### Image analysis and quantification

Maximum intensity projections were created from acquired image z-stacks using ZEN-Blue software (Zeiss) or FIJI (Schindelin et al., 2012). All confocal images and analyses are based on maximum intensity projections of the entire z-stack, except for Figure 4A-C, Figure 4–figure supplement 1A, and Figure 6F and G, which are maximum intensity projections of 1.8-7.6 µm in the apical junctional domain to highlight junctional myosin IIB signal. Drift that occurred while acquiring z-stacks of GFP-Vinculin in live embryos was corrected using the StackReg plugin in FIJI (Thévenaz et al., 1998).

ZO-1, myosin IIB, and pMRLC planar polarity measurements (Figure 1C and D) were measured in 50 µm x 50 µm regions using SIESTA software (Fernandez-Gonzalez and Zallen, 2011). Planar polarity was calculated by dividing the mean intensity of ML junctions (oriented at 0-15° relative to the mediolateral axis) by the mean intensity of AP junctions (oriented at 75-90° relative to the mediolateral axis) after subtracting background signal. Background signal was calculated as the average of the mean pixel intensities in 20-40 circular cytoplasmic regions drawn in FIJI.

Apical cell areas (Figure 2D-F, Figure 3D-H, Figure 7—figure supplement 1A and B) were measured in a 100 µm x 100 µm lateral region of the midbrain. N-cadherin was used to label the apical borders of cells in early and late elevation embryos and ZO-1 was used to label the apical borders of cells in pre-elevation embryos, as N-cadherin expression is low at pre-elevation stages (Brooks et al., 2025). GFP-Plekha7 was used to label the apical adherens junctions in time-lapse movies. Cell segmentation was performed using SeedWater Segmenter (Mashburn et al., 2012), and apical cell areas were quantified and heat maps were generated using the MorphLibJ plugin in FIJI (Legland et al., 2016).

The apical and basal spans of the midbrain neuroepithelium (Figure 3C) were measured by manually drawing lines along the apical and basal borders of the neural plate from the non-neural ectoderm-neuroepithelial border of one neural fold to the other using the segmented line tool in FIJI. Cell height (Figure 3–Supplemental Figure 2A) was determined by measuring the distance between the apical and basal borders of the neural plate, roughly halfway between the non-neural ectoderm-neuroepithelial border and the midline for each neural fold.

The percentage of mitotic cells (Figure 3—figure supplement 2B and C) was calculated by dividing the number of phospho-histone H3-positive nuclei by the total number of cells in a 100 µm x 100 µm region of the midbrain in 7-9 somite wild-type and *Vinculin^ΔEpi^* embryos, using ZO-1 or GFP-Plekha7 to visualize apical cell outlines. Total cell counts were determined by segmenting cells using SeedWater Segmenter (Mashburn et al., 2012).

GFP-Vinculin intensity at tricellular junctions (TCJ intensity) was measured in FIJI as the mean intensity of a circular region of interest (ROI) centered on the tricellular junction (Figure 4K). The GFP-Vinculin and GFP-Plekha7 TCJ ratios (Figure 4H and I) were calculated in FIJI by dividing by dividing the TCJ intensity by the average mean intensities of lines drawn along the three connected bicellular junctions after subtracting background signal as calculated above.

Intensity profiles of the width of the myosin IIB and F-actin signal at bicellular junctions (Figure 4—figure supplement 1C and D) were measured along 3 µm-long lines perpendicular to the bicellular junction and centered on the myosin IIB signal in FIJI. An average value was obtained for 10 bicellular junctions/neural fold after subtracting background signal as calculated above and normalized to the maximum intensity for each line (8 neural folds were analyzed in 4 embryos).

Western blots (Figure 4—figure supplement 3A-C) were quantified by measuring the mean grey value of a box drawn around each band. The same sized box was used for each band of a particular protein and band intensities were normalized to the intensity of the β-catenin loading control.

The number of adherens junction and tight junction gaps (Figure 5E and F, Figure 6B and C, Figure 5— figure supplement 1G, Figure 6—figure supplement 1B and C) was quantified by manually counting regions of aberrant loss of junctional signal in 50 µm x 50 µm regions of the lateral midbrain. The percentage of 3, 4, and 5+ cell junctions with gaps (Figure 5G-I) was determined by dividing the number of gaps at each type of junction by the total number of that type of junction in each image. The areas of adherens junction and tight junction gaps (Figure 6B-E, Figure 6—figure supplement 1B and C) were determined by drawing circular ROIs over gaps in signal, with the diameter of the ROI representing the furthest points of lost signal. Places where ROIs for multiple gaps overlapped were treated as one large gap and a new circular ROI was redrawn to cover the overlapping gaps. The average gap area at tight junctions was measured at each site in which there was an adherens junction gap, and scored as a value of 0 if no corresponding tight junction gap was detected. Of the adherens junction gaps observed, 141/150 were not associated with tight junction gaps in Control^ESC^ and 442/498 were not associated with tight junction gaps in *Vinculin^ESC^*. Gaps at bicellular junctions (plotted in Figure 5—figure supplement 1A and B) were not included in the other adherens junction gap measurements in Figure 5, Figure 6, Figure 5—figure supplement 1G, and Figure 6—figure supplement 1.

Intensity profiles along orthogonal XZ reslices of AiryScan z-stacks at bicellular junctions (Figure 6—figure supplement 1D and E) were measured along 3 µm-long lines drawn from apical to basal and centered on the ZO-1 signal. An average value was obtained for 10 bicellular junctions/neural fold and normalized to the maximum intensity for each line and (4 neural folds/embryo were analyzed in 2 embryos).

For analysis of time-lapse movies (Figure 7, Figure 7—figure supplement 1), maximum intensity projections were registered in XY using the Correct 3D drift plugin in FIJI. Cell behavior was analyzed in 100 µm x 100 µm regions in the lateral midbrain. Junctions where 5 or more cells meet were scored as defective when there was a clear loss of GFP-Plekha7 signal at the central high-order vertex. Dividing cells were identified by the presence an ingressing cleavage furrow, which generates a distinctive figure-8 morphology, and t=0 was defined as one time point before the cleavage furrow started to ingress. Dividing cells were tracked every 6 min until a clear GFP-Plekha7-positive 4-cell vertex, a new interface between daughter cells, or a new interface between neighboring cells that separated the two daughter cells was detected. Cell divisions were scored as having a gap in GFP-Plekha7 signal at neighbor contacts if a significant reduction in GFP-Plekha7 signal was detected at the interface between the dividing cell and surrounding cells that was not restored within 90 minutes (Figure 7E, left). Cell divisions were scored as having a gap in GFP-Plekha7 signal at new contacts if GFP-Plekha7 was not detected at a new vertex or interface within 90 minutes after the onset of ingression (Figure 7E, right). Only cell divisions that began within the first 90 minutes of the 3-hour movie were analyzed.

### Statistics and figure preparation

Statistical analysis and graph preparation were performed with GraphPad Prism. Statistical analyses used were the unpaired t-test with Welch’s correction, which does not assume equal standard deviations, and the Kolmogorov-Smirnov test to compare pooled distributions. See Supplementary File 1 for all n and p values. Figures were assembled using Adobe Illustrator.

## Acknowledgments

The authors thank Corey Elowsky, Kazi Hossain, Lila Neahring, Carlos Patiño Descovich, Nathan Shugarts-Devanapally, Masako Tamada, and Carolyn Ton for helpful feedback on the manuscript. We are grateful to LARGE, RIKEN BDR (Kobe, Japan) for technical advice and provision of the HK3i ESC line, the *R26-H2B-EGFP* mouse line (Abe et al., 2011; RIKEN accession No. CDB0238K), and the *R26-PHA7-EGFP* mouse line (Shioi et al., 2017) (RIKEN accession No. CDB0261K). Use of services by MSK Core Facilities, including animal housing and veterinary services, mouse genome engineering, and DNA sequencing, was supported by NIH/NCI Cancer Center Support Grant P30 CA008748. ERB was supported by NIH/NINDS F32 fellowship NS098832 and startup funds from North Carolina State University. JAZ is an investigator of the Howard Hughes Medical Institute.

## Author contributions

Ian S Prudhomme – Conceptualization, Formal analysis, Investigation, Methodology, Visualization, Writing – original draft, Writing – review and editing; Eric R Brooks – Conceptualization, Investigation, Methodology, Writing – review and editing; Nilay Taneja – Formal analysis, Investigation, Methodology, Visualization, Writing – review and editing; Bhaswati Bhattacharya – Investigation, Methodology; Brian J LaFleche – Methodology; Yas Furuta – Conceptualization and optimization of ESC-based methodology, Writing – review and editing; Jennifer A Zallen – Conceptualization, Formal analysis, Funding acquisition, Supervision, Writing – review and editing.

## Ethics

All animal experiments were conducted in accordance with PHS guidelines and the NIH Guide for the Care and Use of Laboratory Animals and approved Institutional Animal Care and Use Committee protocols (15-08-013 and 90-12-033) of Memorial Sloan Kettering Cancer Center.

---

**Figure 1—figure supplement 1.**
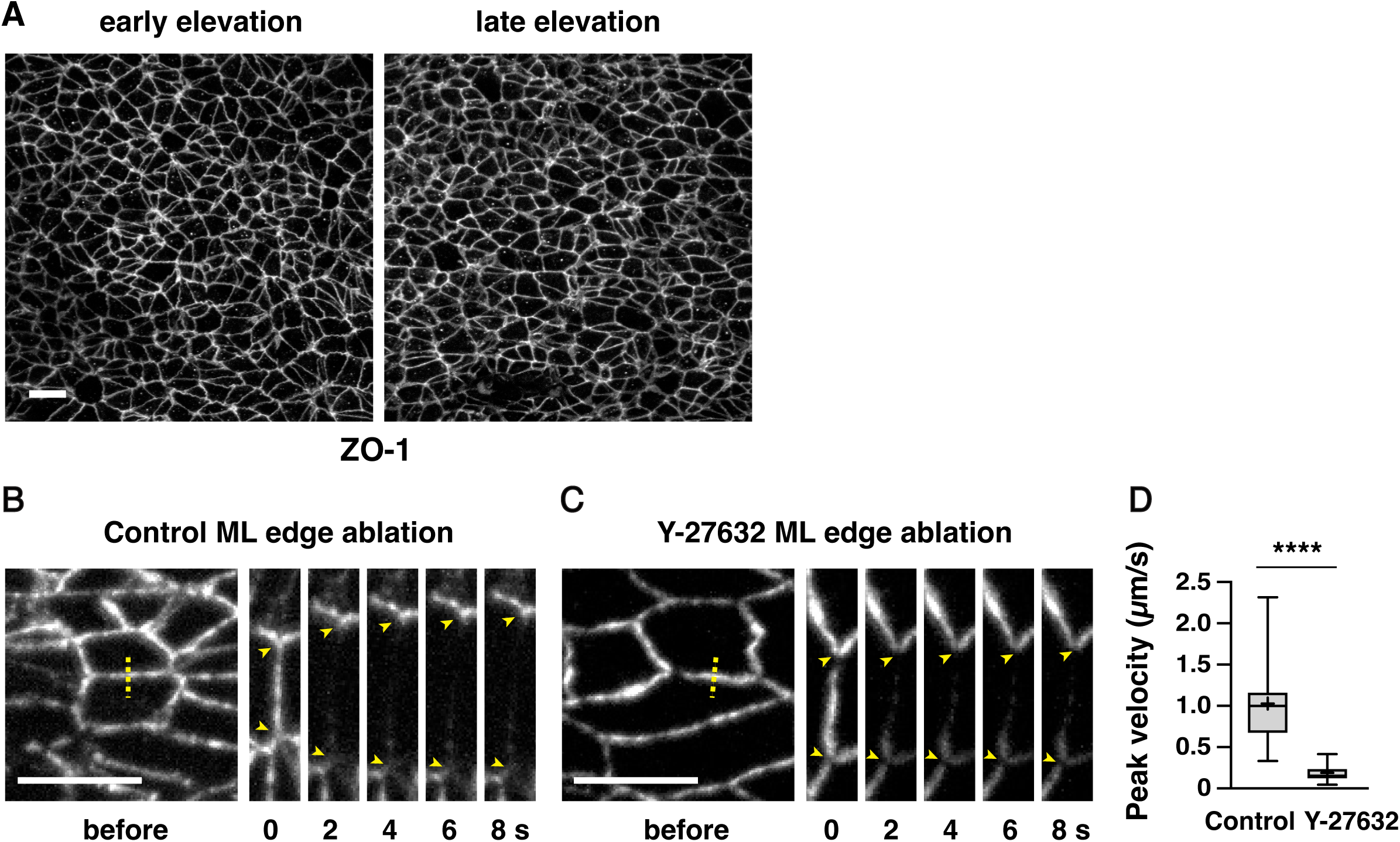
The response to edge ablations in the cranial neural plate requires actomyosin contractility. (A) Localization of the tight junction protein ZO-1 in the lateral midbrain of early and late elevation wild-type embryos (see Figure 1C and D for quantification). (B and C) ML edges before and 2-8 s after ablation in late elevation embryos expressing GFP-Plekha7 and treated for 2 h with water (B) or 200 µM of the Rho-kinase inhibitor Y-27632 (C). (D) Peak recoil velocity after laser ablation of ML edges. Boxes, 25^th^-75^th^ percentile; whiskers, 5^th^-95^th^ percentile; horizontal line, median; +, mean. 15 ablations in 4 embryos/condition. ****p<0.0001 (Welch’s t-test). Maximum intensity projections, anterior up, edges oriented vertically in kymographs. Bars, 10 µm.

**Figure 2—figure supplement 1.**
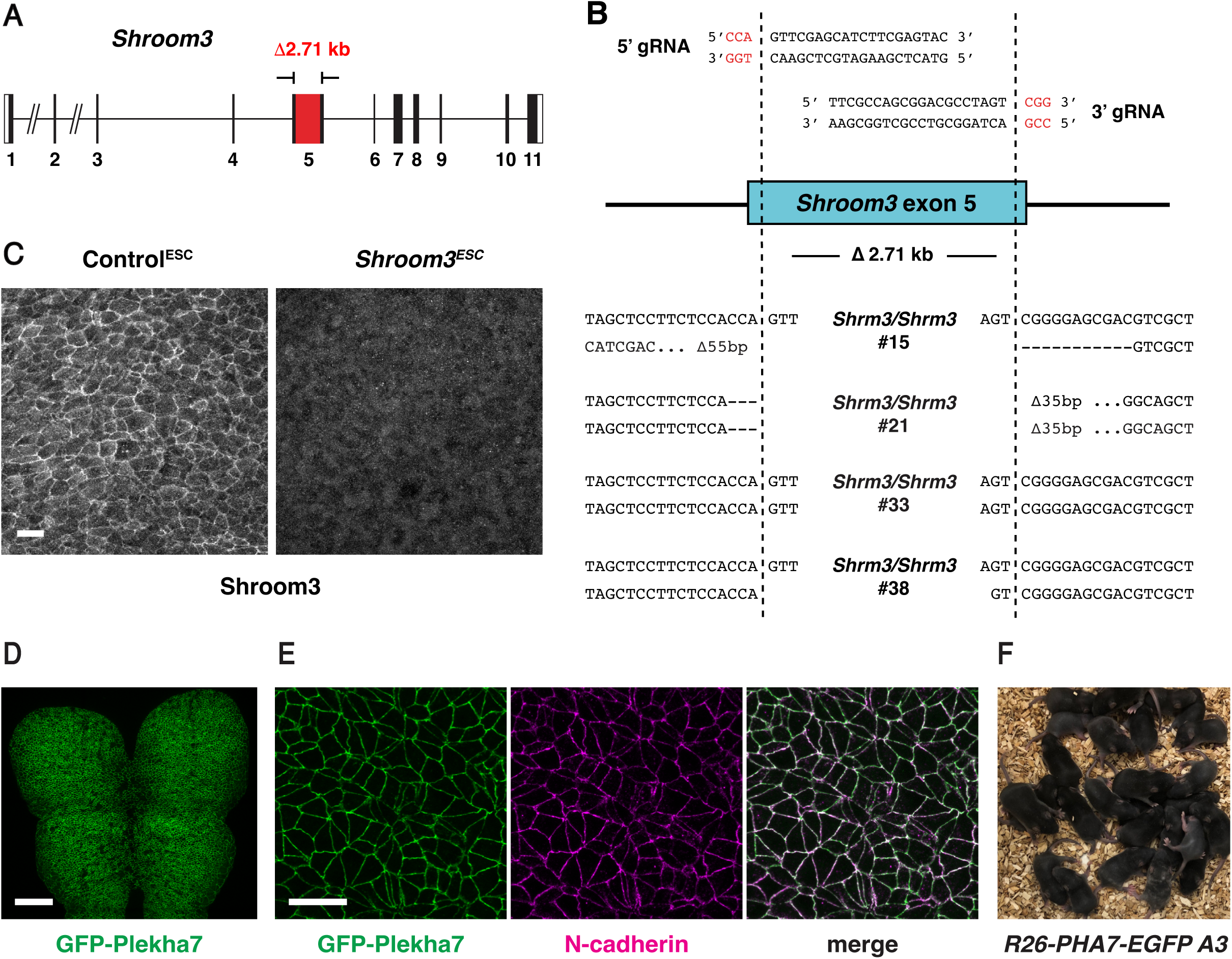
Generation and validation of *Shroom3* mutant and GFP-Plekha7 ESCs. (A and B) ESC clones homozygous for a null mutation in *Shroom3* (A) were identified by PCR screening and deep sequencing (B). (A) Red box, deleted region within exon 5 of the *Shroom3-203* isoform. (B) Deletion breakpoints of four homozygous mutant clones. Top, 5’ and 3’ gRNA target sites. Red, PAM site. Dotted lines, predicted CRISPR cut sites. Bottom, deletion breakpoints. (C) Shroom3 protein is not detected by immunofluorescence in the lateral midbrain in *Shroom3^ESC^* embryos. (D and E) Embryos generated by injection of GFP-Plekha7-expressing ESCs displayed GFP-Plekha7 expression throughout the cranial neural plate (D) (black regions, dividing cells) that colocalized with N-cadherin (E). Maximum intensity projections, anterior up. Bars, 100 μm (D), 10 µm (C and E). (F) Pups generated from the GFP-Plekha7-expressing ESC clone *R26-PHA7-EGFP A3*. 25/27 animals had 100% ESC coat color and 2/2 adults tested displayed germline transmission.

**Figure 3—figure supplement 1.**
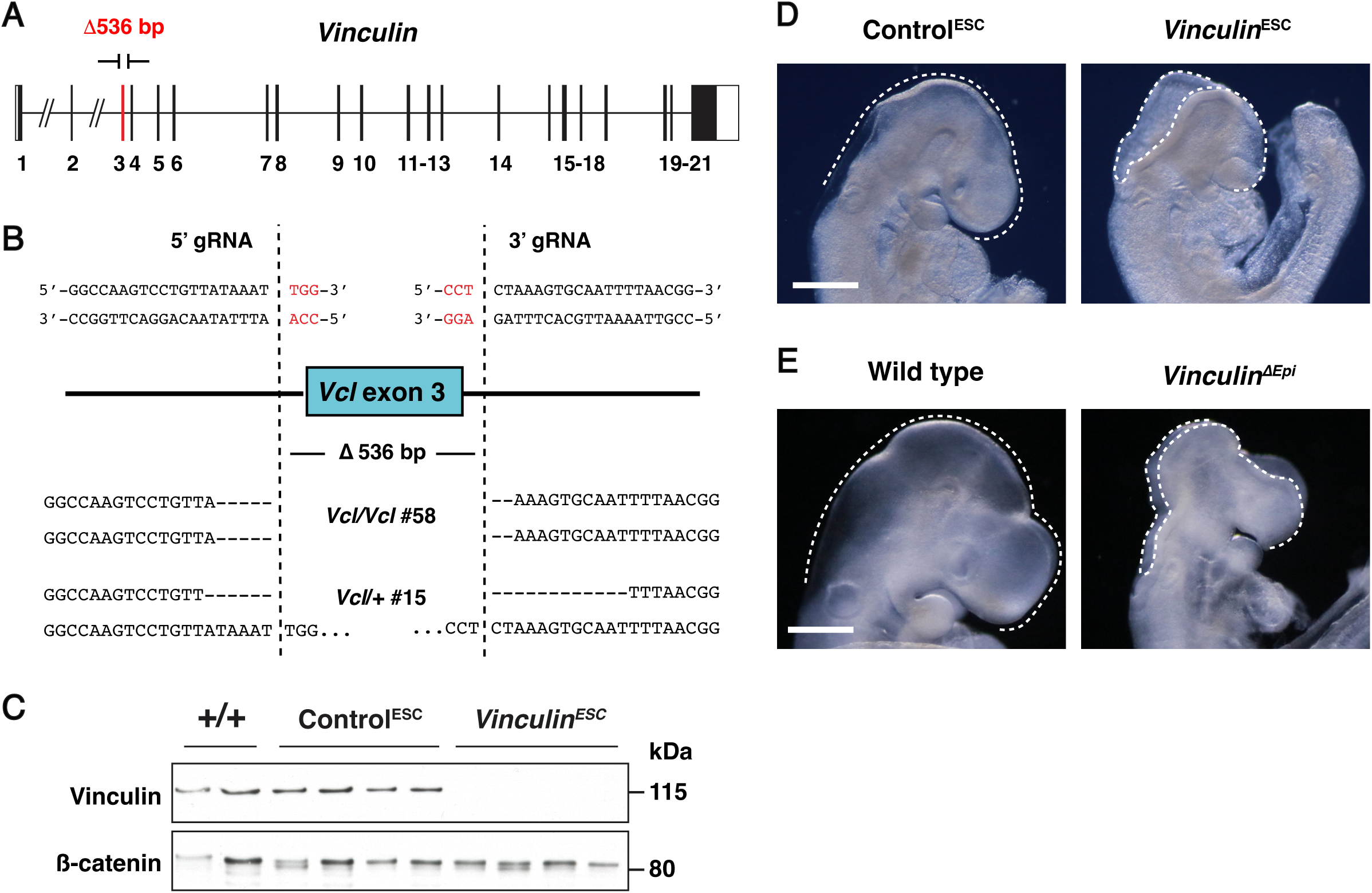
Generation and validation of *Vinculin* mutant ESCs. (A and B) ESC clones homozygous or heterozygous for a null mutation in *Vinculin* (A) were identified by PCR screening and deep sequencing (B). (A) Red box, deleted region including exon 3 of the *Vinculin-201* isoform. (B) Deletion breakpoints of *Vinculin* homozygous mutant clone 58 and heterozygous control clone 15. Top, 5’ and 3’ gRNA target sites. Red, PAM site. Dotted lines, predicted CRISPR cut sites. Bottom, deletion breakpoints. (C) Vinculin protein is absent in *Vinculin^ESC^* embryos (generated from clone 58) and present in Control^ESC^ embryos (generated from clone 15) and +/+ embryos (FVB/N) (one E9.5 embryo/lane). (D) Light micrographs of E9.5 Control^ESC^ and *Vinculin^ESC^* embryos (23/23 *Vinculin^ESC^* embryos and 0/15 Control^ESC^ embryos displayed exencephaly). (E) Light micrographs of E9.5 wild-type and *Vinculin^ΔEpi^* embryos (11/11 *Vinculin^ΔEpi^* embryos and 0/30 wild-type and heterozygous littermate controls displayed exencephaly). Lateral views, dotted lines indicate the lateral edges of the cranial neural plate. Bars, 500 µm.

**Figure 3—figure supplement 2.**
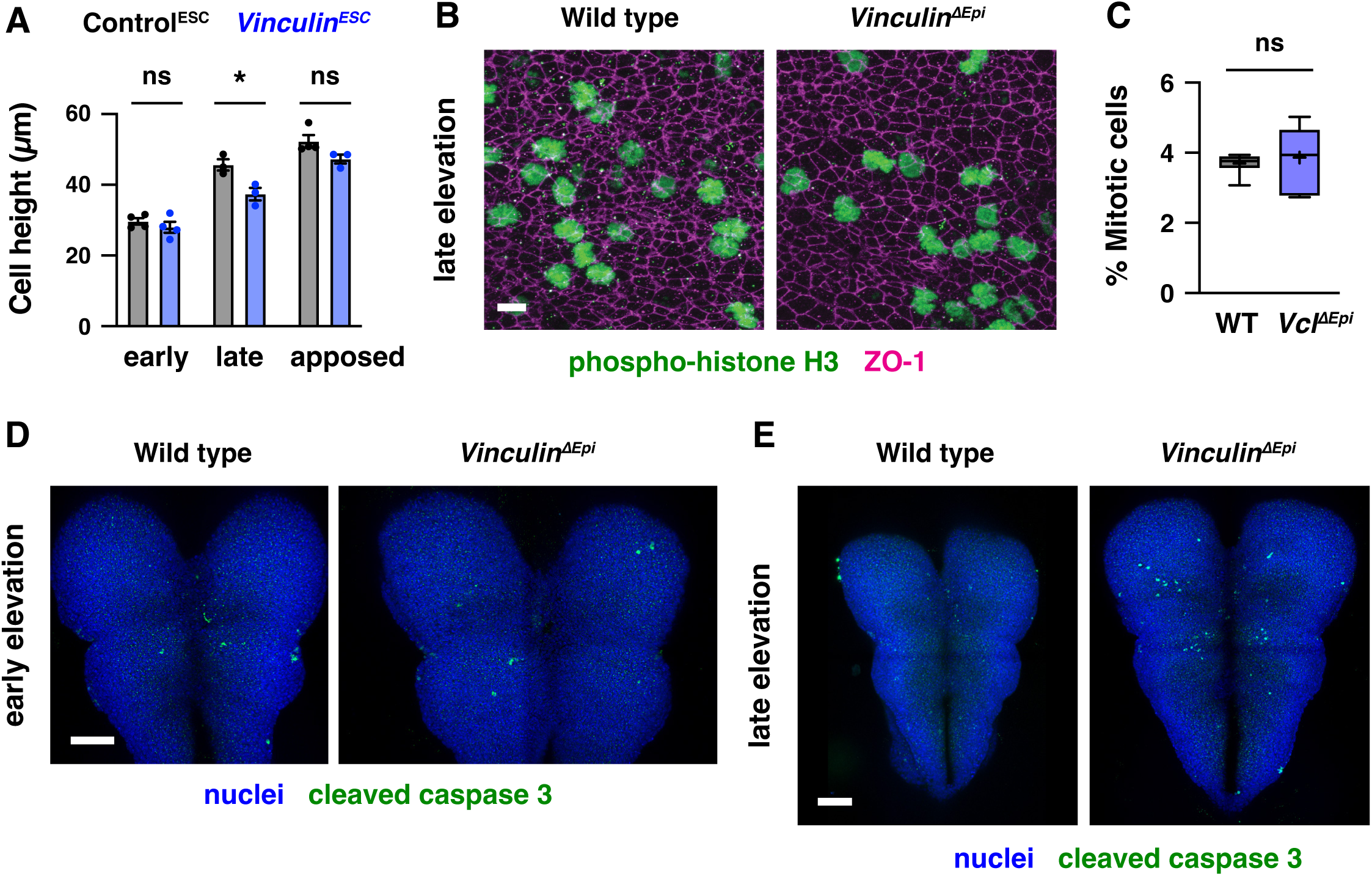
Apical-basal elongation, proliferation, and apoptosis are unaffected in *Vinculin* mutants. (A) Apical-basal cell height in transverse sections of the lateral midbrain of Control^ESC^ and *Vinculin^ESC^*embryos in early elevation (4-6 somites), late elevation (7-9 somites), and apposition (8-10 somites). A single value was obtained for each embryo and the mean±SEM between embryos is shown (3-4 embryos/genotype). *p<0.03 (Welch’s t-test). (B) Dividing cells detected with phospho-histone H3 in the lateral midbrain of wild-type and *Vinculin^ΔEpi^*embryos in late elevation. Cell outlines are visualized with ZO-1. (C) Percentage of cells positive for phospho-histone H3 in wild-type and *Vinculin^ΔEpi^* embryos. Boxes, 25^th^-75^th^ percentile; whiskers, 5^th^-95^th^ percentile; horizontal line, median; +, mean, 6-7 regions in 3-4 embryos/genotype. (D, E) Detection of the apoptotic cell marker cleaved caspase 3 is similar in wild-type and *Vinculin^ΔEpi^* embryos in early (D) and late (E) elevation. Maximum intensity projections, anterior up. Bars, 10 µm (B), 100 µm (D, E).

**Figure 4—figure supplement 1.**
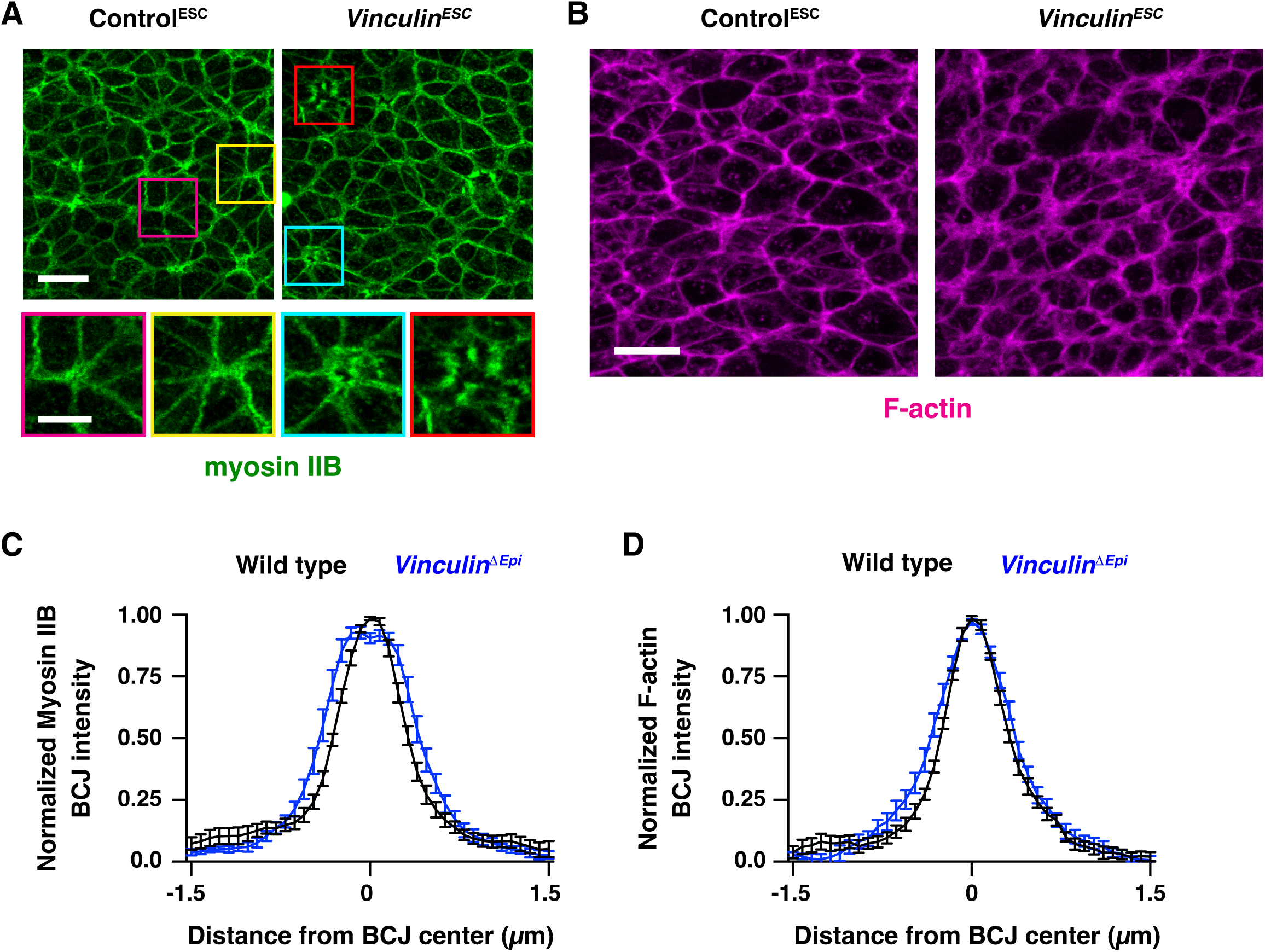
Actomyosin localization in *Vinculin^ESC^* and *Vinculin^ΔEpi^* embryos. (A) Localization of myosin IIB in Control^ESC^ and *Vinculin^ESC^* embryos in late elevation. Bottom panels, examples of multicellular junctions with wild-type myosin IIB localization (left) and moderate (cyan) or severe (red) gaps in myosin II localization (right). (B) Localization of F-actin (phalloidin) in Control^ESC^ and *Vinculin^ESC^* embryos in late elevation. (C and D) Normalized intensity profiles of myosin IIB (C) and F-actin (phalloidin) (D) along 3 μm lines perpendicular to bicellular junctions. An average value was obtained for 10 bicellular junctions/neural fold and the mean±SEM between neural folds is shown (8 neural folds in 4 embryos/genotype). Maximum intensity projections, anterior up. Bars, 10 µm (A, top panels, B), 2 µm (A, bottom panels).

**Figure 4—figure supplement 2.**
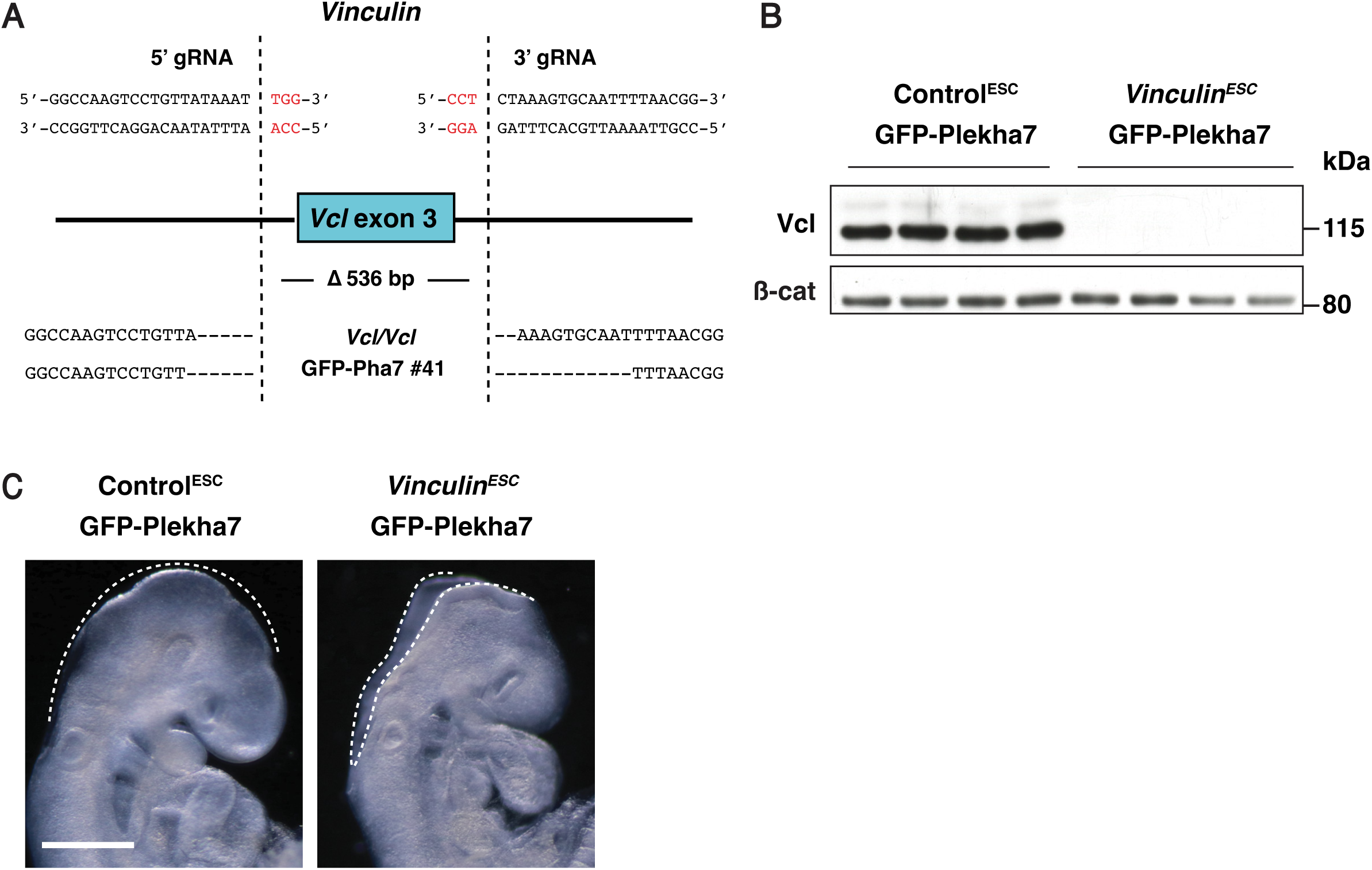
Generation and validation of *Vinculin* mutant ESCs expressing GFP-Plekha7. (A) Deletion breakpoints of *Vinculin* homozygous mutant clone 41 expressing GFP-Plekha7. Top, 5’ and 3’ gRNA target sites. Red, PAM site. Dotted lines, predicted CRISPR cut sites. Bottom, deletion breakpoints based on deep sequencing of amplicons around the 5’ and 3’ cut sites. (B) Vinculin protein is absent in *Vinculin^ESC^*GFP-Plekha7 embryos (generated from clone 41) and present in Control^ESC^ GFP-Plekha7 embryos (generated from unedited wild-type GFP-Plekha7 ESCs) (one E9.5 embryo/lane). (C) Light micrographs of E9.5 Control^ESC^ GFP-Plekha7 and *Vinculin^ESC^* GFP-Plekha7 embryos (28/28 *Vinculin^ESC^* GFP-Plekha7 embryos and 0/30 Control^ESC^ GFP-Plekha7 embryos displayed exencephaly). Lateral views, dotted lines indicate the lateral edges of the cranial neural plate. Bar, 500 µm.

**Figure 4—figure supplement 3.**
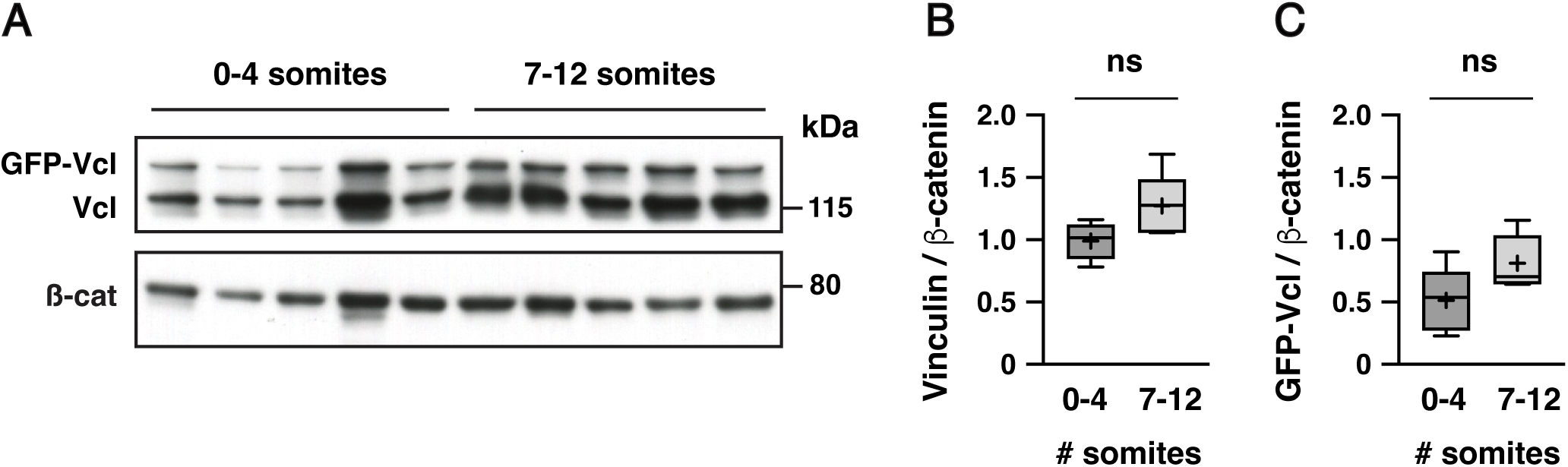
Quantification of GFP-Vinculin expression from the *R26* locus in ESC-derived embryos. (A) Western blot showing the presence of GFP-Vinculin in addition to the endogenous Vinculin protein detected with the anti-Vinculin antibody (one embryo/lane). (B, C) Vinculin (B) and GFP-Vinculin (C) proteins showed no significant difference in total protein levels between 0-4 somite and 7-12 somite stages. Protein intensity was normalized to the intensity of the β-catenin loading control. Boxes, 25^th^-75^th^ percentile; whiskers, 5^th^-95^th^ percentile; horizontal line, median; +, mean.

**Figure 5—figure supplement 1.**
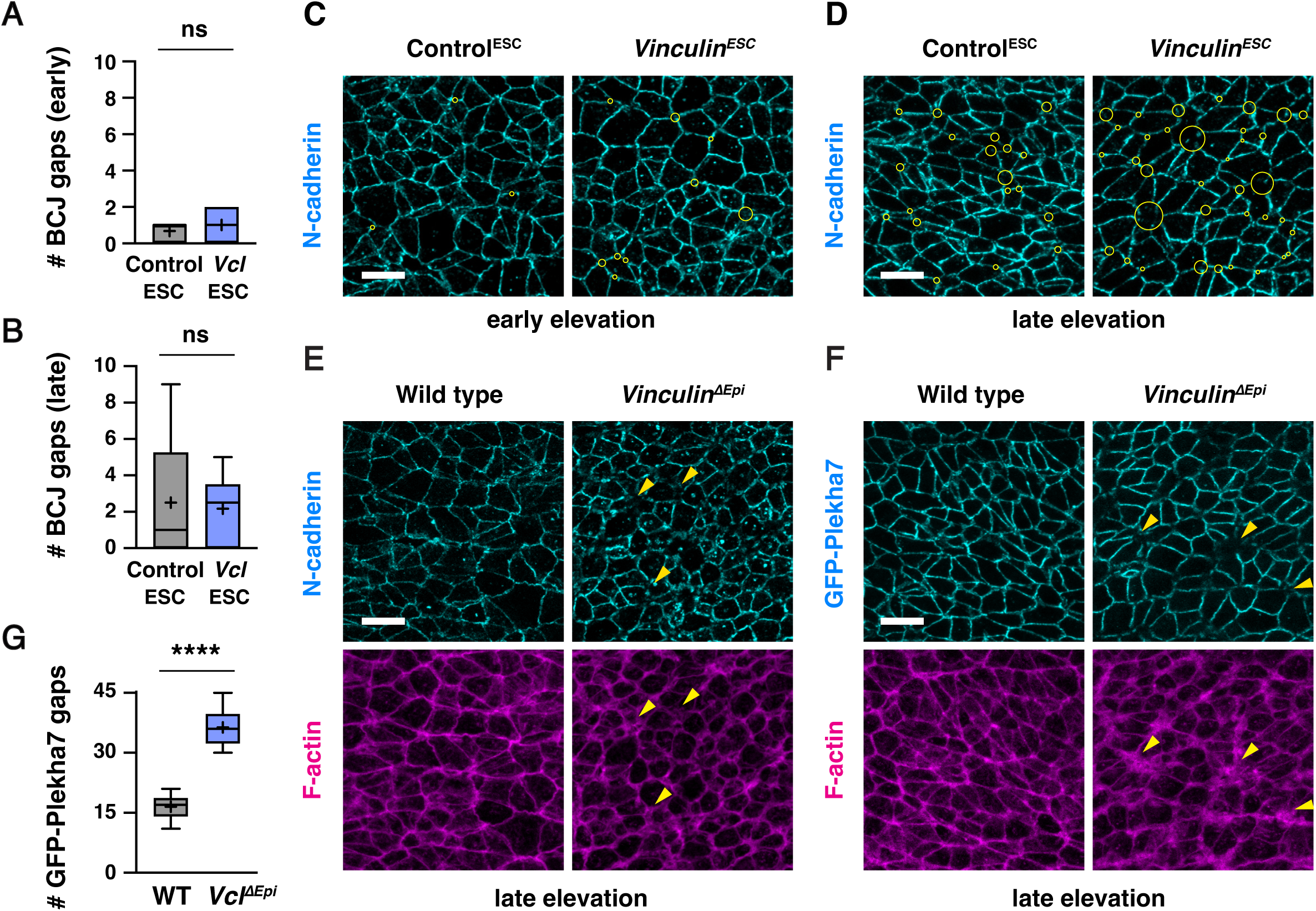
*Vinculin^ΔEpi^* embryos display defects in F-actin and adherens junction localization. (A and B) Number of gaps in N-cadherin localization at bicellular junctions in a 50 µm x 50 µm region of the lateral midbrain in Control^ESC^ and *Vinculin^ESC^*embryos in early (A) and late (B) elevation. (C, D) Localization of N-cadherin in Control^ESC^ and *Vinculin^ESC^* embryos in early and late elevation (reproduced from Figure 5A and B). Yellow circles show all tricellular and multicellular junctions scored as defective. (E) Localization of N-cadherin and F-actin (phalloidin) in wild-type and *Vinculin^ΔEpi^* embryos in late elevation. (F and G) Localization of GFP-Plekha7 and F-actin (phalloidin) (F) and number of gaps in GFP-Pha7 localization (G) in a 50 µm x 50 µm region of the lateral midbrain in wild-type and *Vinculin^ΔEpi^* embryos in late elevation. Arrowheads show examples of multicellular junctions scored as defective. 6 regions in 3 embryos/genotype in A, B, and G. Maximum intensity projections, anterior up. Bars, 10 µm.

**Figure 6—figure supplement 1.**
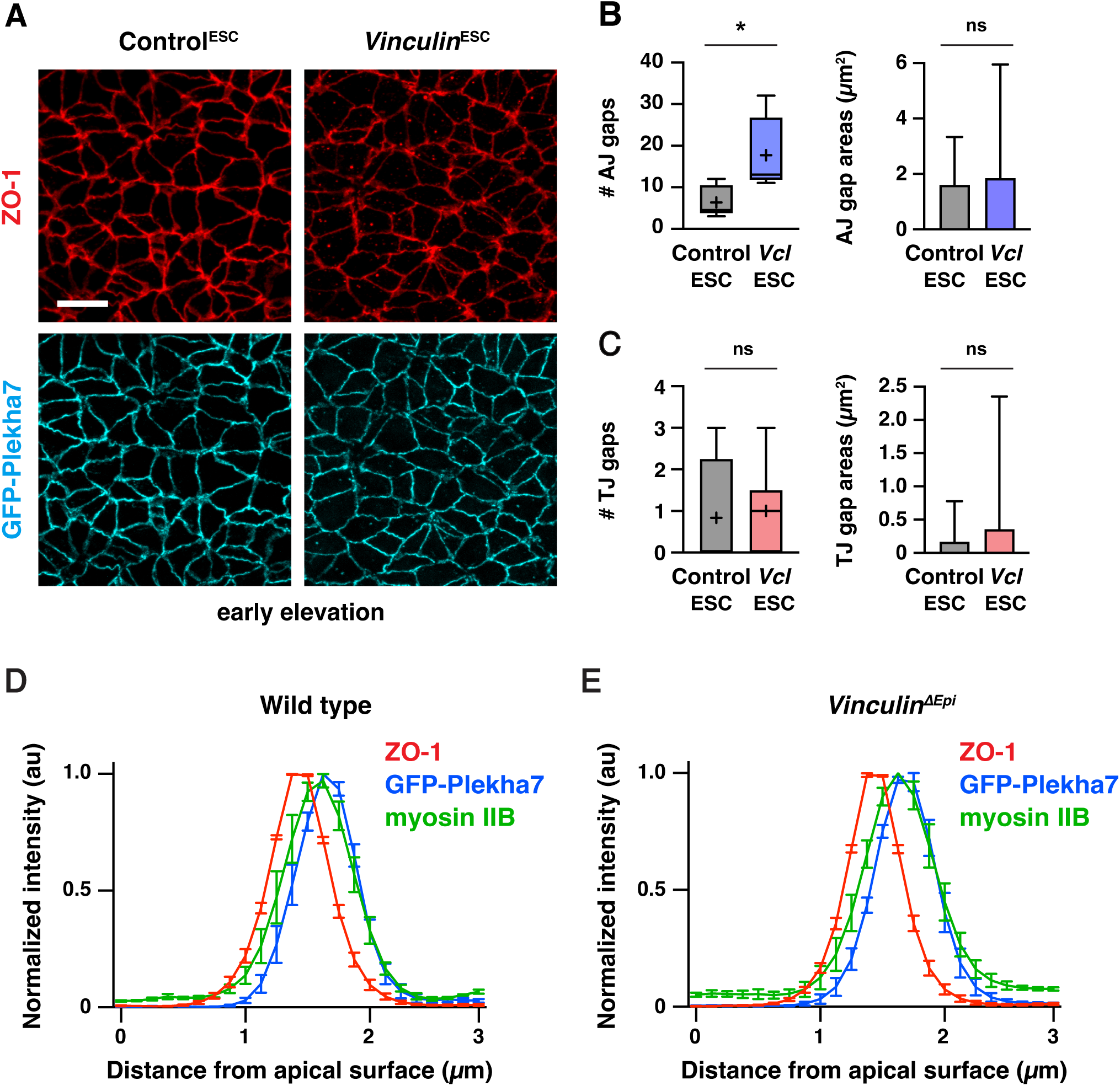
Tight junction and adherens junction protein localization in *Vinculin^ESC^* and *Vinculin^ΔEpi^* embryos. (A) Localization of the tight junction protein ZO-1 and the adherens junction protein GFP-Plekha7 in early elevation Control^ESC^ and *Vinculin^ESC^* embryos. (B, C) Number (left) and areas (right) of adherens junction gaps (B) and tight junction gaps (C) in a 50 µm x 50 µm region in early elevation Control^ESC^ and *Vinculin^ESC^* embryos. Note the differences in scale between the adherens junction and tight junction plots. Boxes, 25^th^-75^th^ percentile; whiskers, 5^th^-95^th^ percentile; horizontal line, median; +, mean. 38-106 gaps from 6 regions in 3 embryos. *p<0.03 (Welch’s t-test). (D and E) Normalized intensity profiles of ZO-1, GFP-Plekha7, and myosin IIB measured along 3 μm lines parallel to the apical-basal axis of bicellular junctions in XZ reconstructions of AiryScan z-stacks in late elevation wild-type and *Vinculin*^Δ*Epi*^ embryos. An average value was obtained for 10 bicellular junctions/neural fold and the mean±SEM between neural folds is shown (4 neural folds in 2 embryos/genotype). Maximum intensity projections, anterior up. Bar, 10 µm.

**Figure 7—figure supplement 1.**
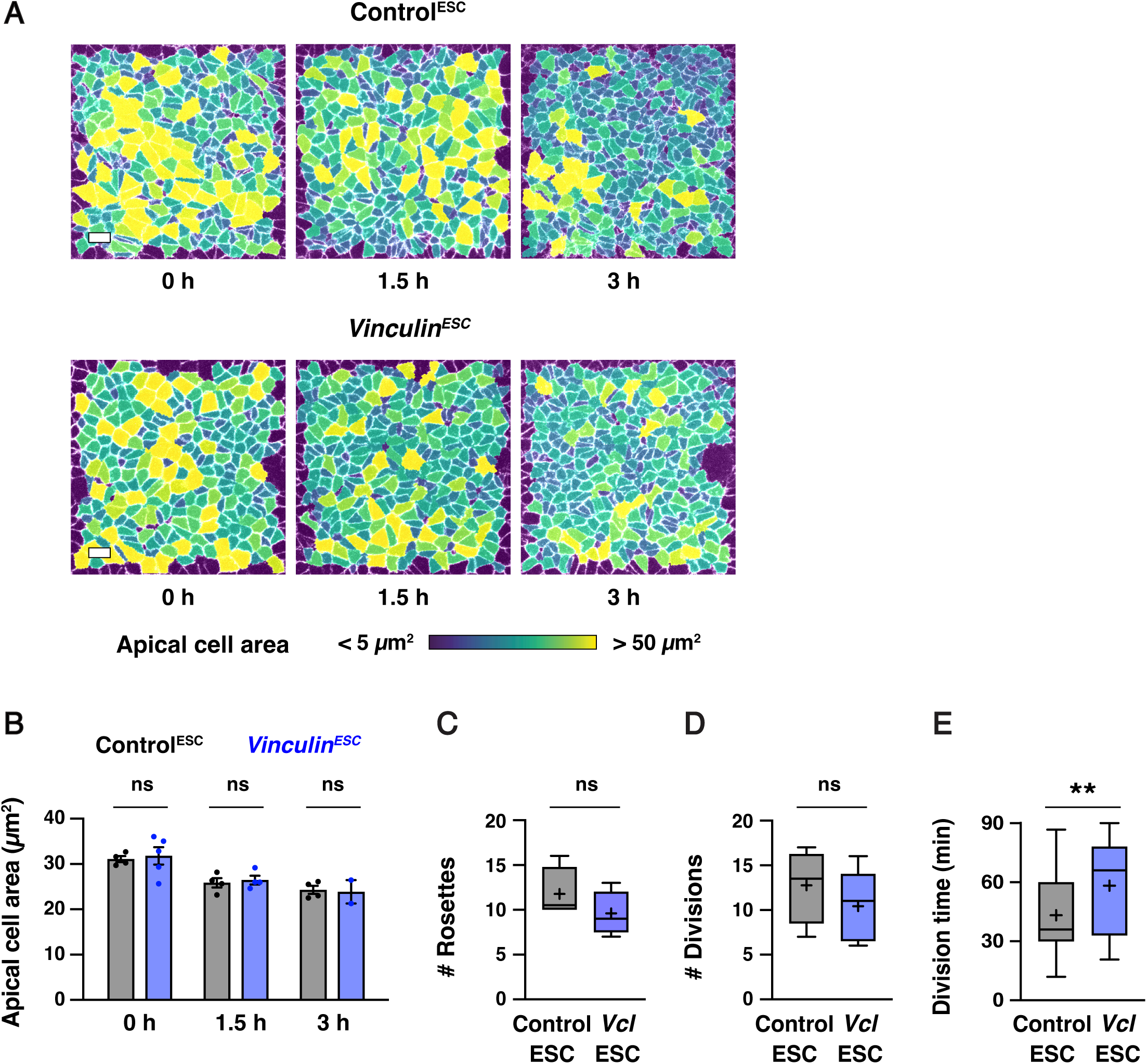
Apical constriction, high-order junctions, and cell division initiate correctly in *Vinculin^ESC^* embryos. (A) Stills from time-lapse movies of Control^ESC^ (top) and *Vinculin^ESC^*(bottom) embryos expressing GFP-Plekha7, color coded by apical cell area. (B) The average apical cell area decreases similarly in Control^ESC^ embryos and in *Vinculin^ESC^* embryos that did not display severe defects in cell adhesion. n=4 Control^ESC^ embryos/time point, 5 *Vinculin^ESC^* embryos at 0 h, 4 *Vinculin^ESC^* embryos at 1.5 h, and 2 *Vinculin^ESC^* embryos at 3.0 h. (C, D) The numbers of rosettes (C) and dividing cells (D) were not significantly different in Control^ESC^and *Vinculin^ESC^* embryos. (E) Division times (time elapsed between the onset of cleavage furrow ingression and the appearance of GFP-Plekha7 at the new vertex or interface) in Control^ESC^ (top) and *Vinculin^ESC^* (bottom) embryos. A single value was obtained for each embryo and the mean±SEM between embryos is shown in (B). Boxes, 25^th^-75^th^ percentile; whiskers, 5^th^-95^th^ percentile; horizontal line, median; +, mean. 4-5 embryos/genotype in (C-E). Maximum intensity projections, anterior up. Bars, 10 µm.

**Supplementary File 1.**
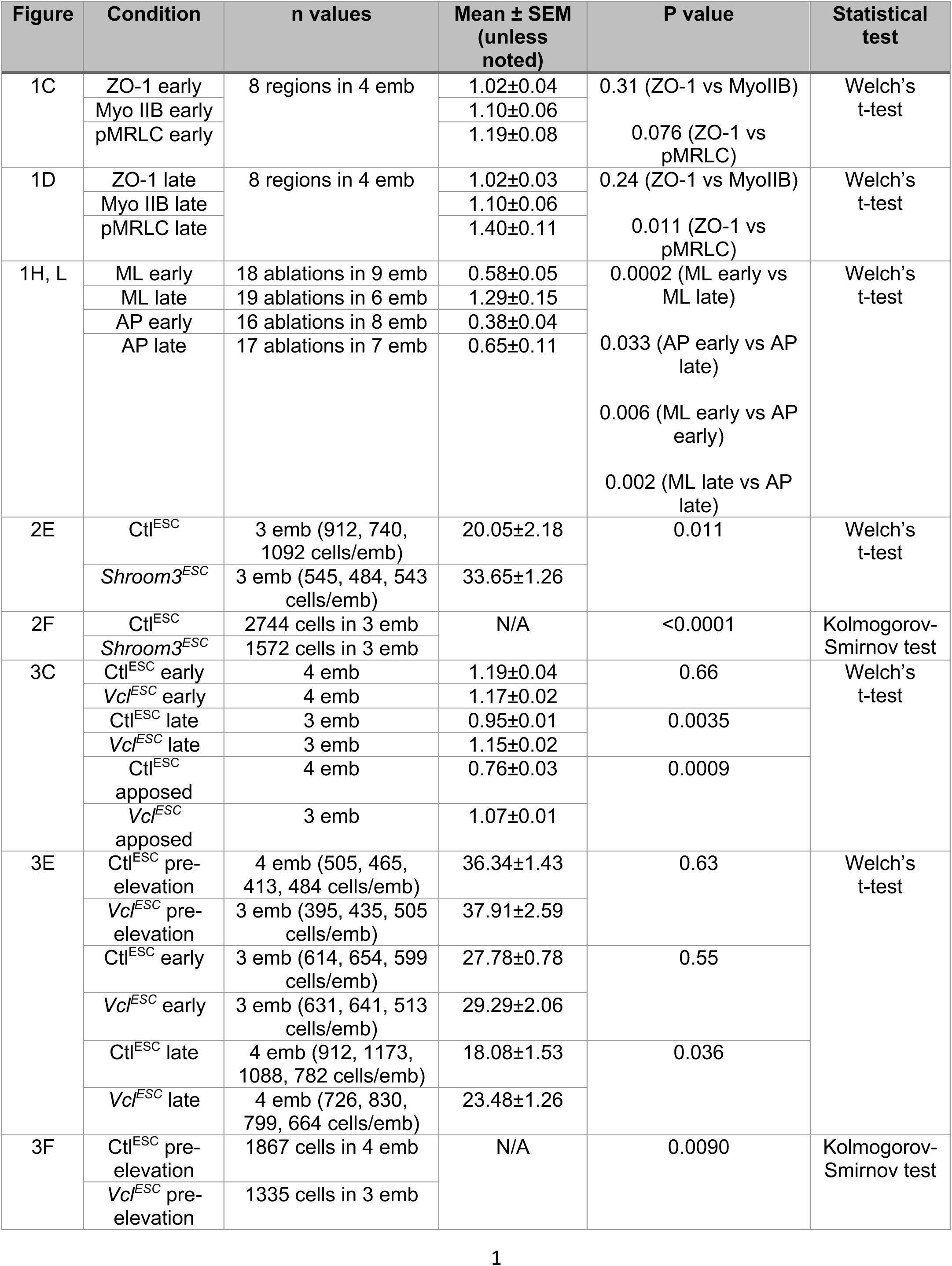
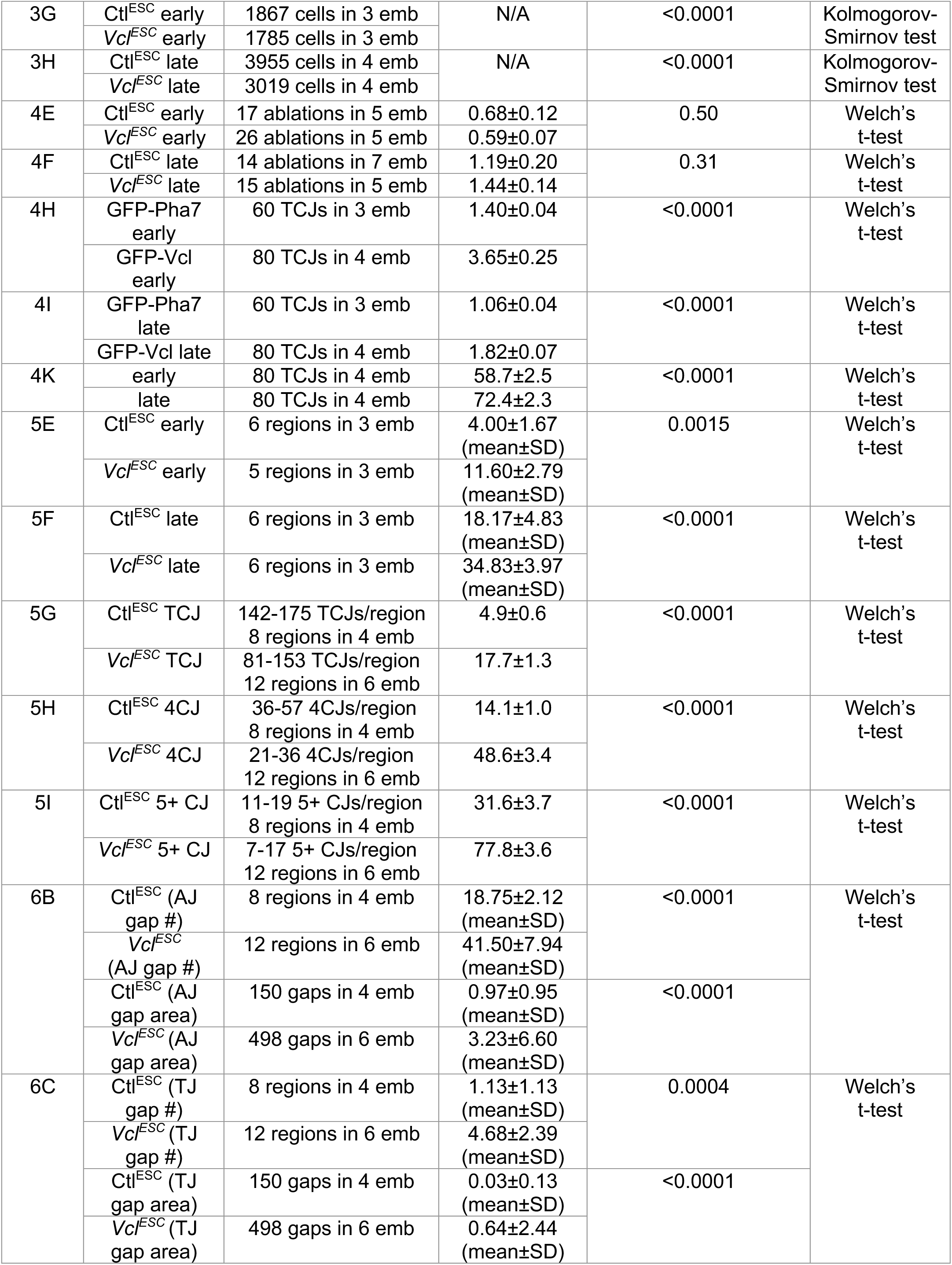
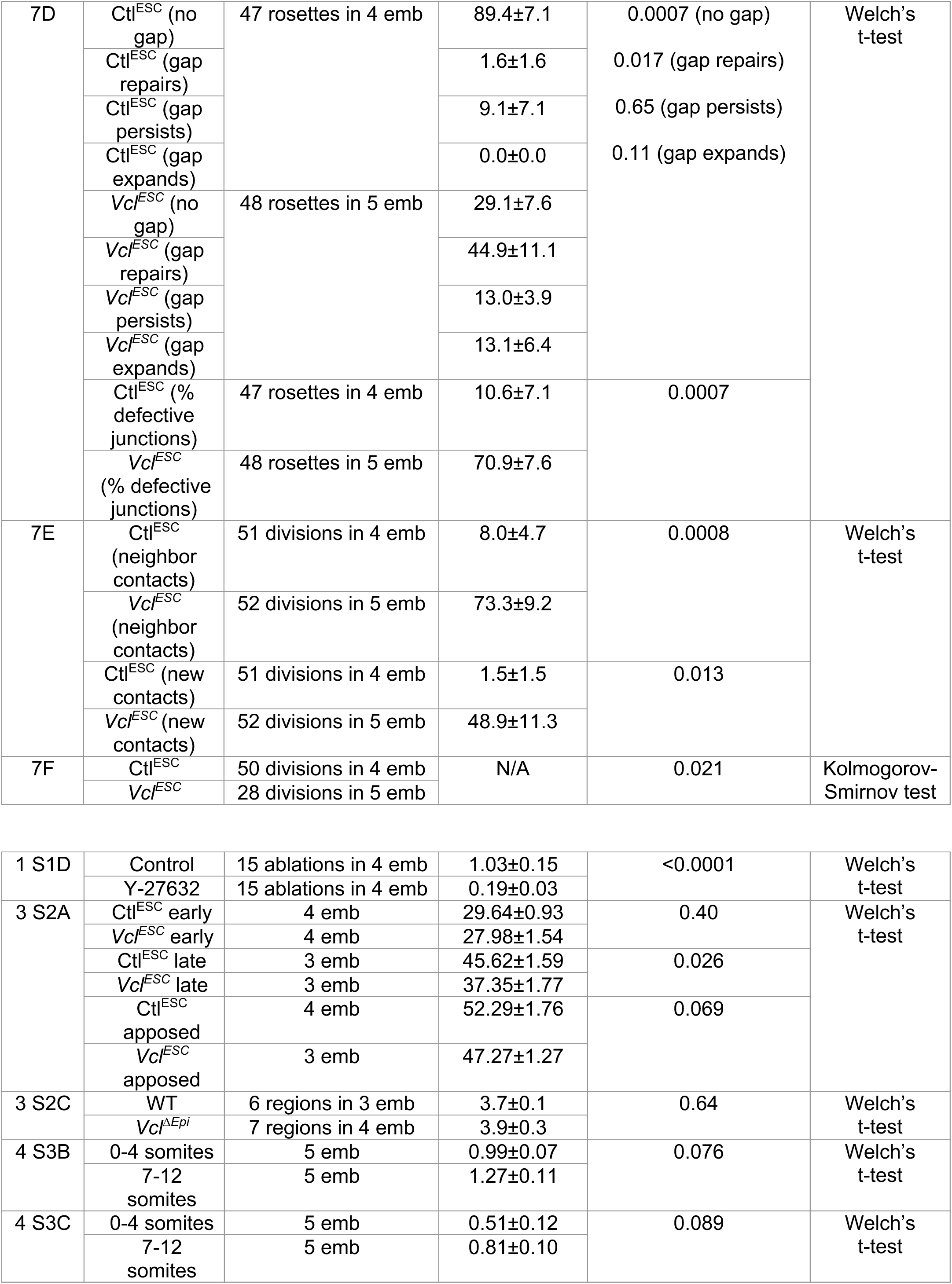
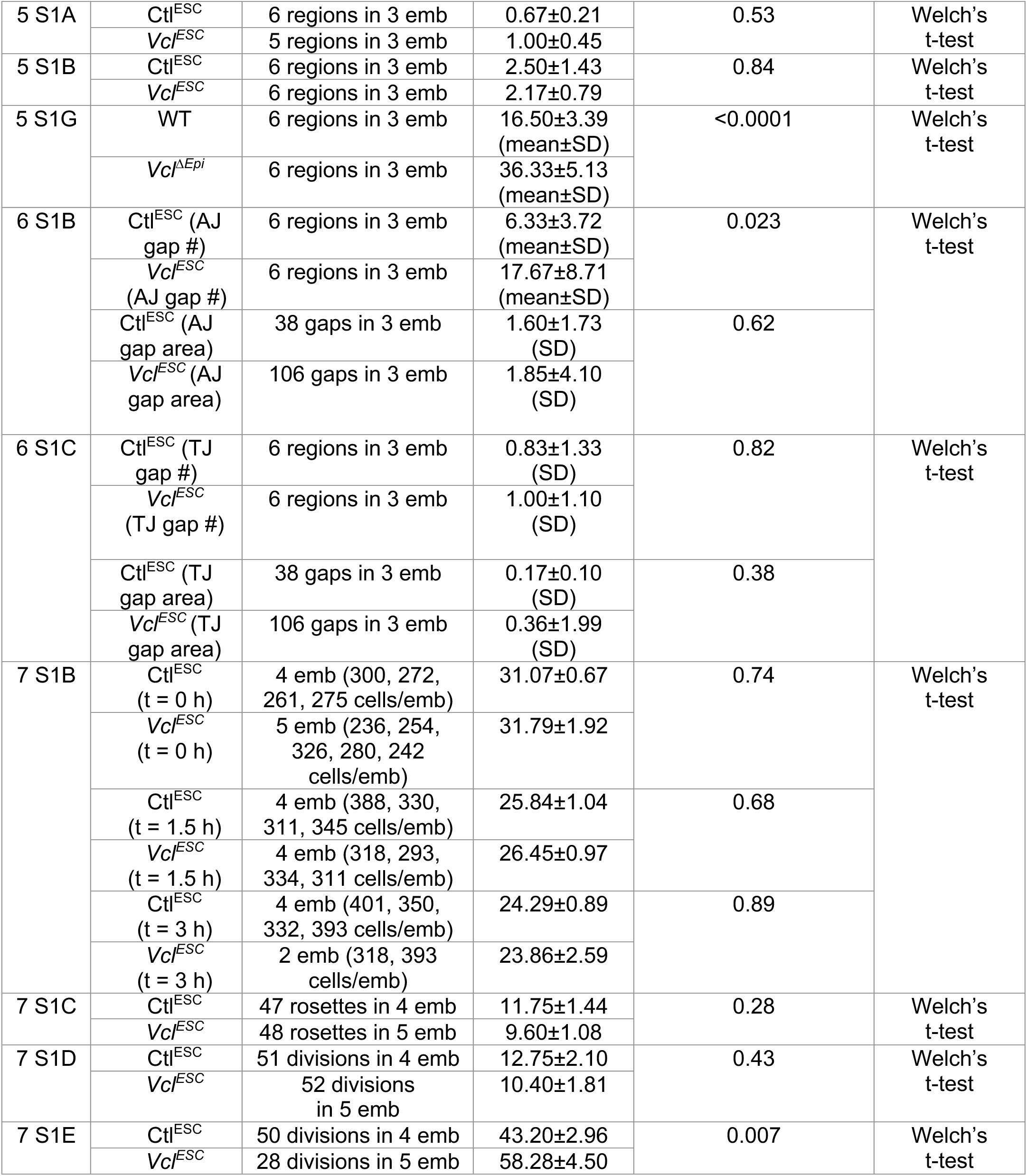
N values and details of statistical analyses performed.

**Supplementary Movie 1. Time-lapse movie of a Control^ESC^ embryo expressing GFP-Plekha7.** Images were acquired every 6 min for 3 hr. Maximum intensity projections, anterior up. Bar, 10 μm.

**Supplementary Movie 2. Time-lapse movie of a moderately defective *Vinculin^ESC^* embryo expressing GFP-Plekha7.** Images were acquired every 6 min for 3 hr. Maximum intensity projections, anterior up. Bar, 10 μm.

**Supplementary Movie 3. Time-lapse movie of a severely defective *Vinculin^ESC^* embryo expressing GFP-Plekha7.** Images were acquired every 6 min for 3 hr. Maximum intensity projections, anterior up. Bar, 10 μm.

